# Single Cell RNA Profiling Reveals Adipocyte to Macrophage Signaling Sufficient to Enhance Thermogenesis

**DOI:** 10.1101/2020.05.04.077529

**Authors:** Felipe Henriques, Alexander H. Bedard, Adilson Guilherme, Mark Kelly, Jingyi Chi, Peng Zhang, Lawrence M. Lifshitz, Karl Bellvé, Leslie Rowland, Batuhan Yenilmez, Shreya Kumar, Yetao Wang, Jeremy Luban, Lee S. Weinstein, Jiandie D. Lin, Paul Cohen, Michael P. Czech

## Abstract

The “browning” of inguinal white adipose tissue (iWAT) through increased abundance of thermogenic beige/brite adipocytes is induced by cold exposure and many other perturbations in association with beneficial systemic metabolic effects. Adipose browning is reported to require activation of sympathetic nerve fibers (SNF), aided by alternately activated macrophages within iWAT. Here we demonstrate the first example of a non-cell autonomous pathway for iWAT browning that is fully independent of SNF activity. Thus, the strong induction of thermogenic adipocytes prompted by deletion of adipocyte fatty acid synthase (iAdFASNKO mice) was unaffected by denervation or the deletion of SNF modulator Neuregulin-4. However, browning of iWAT in iAdFASNKO mice does require adipocyte cAMP/protein kinase A signaling, as it was blocked in adipocyte-selective Fasn/Gsα double KO mice. Single-cell transcriptomic analysis of iAdFASNKO mouse adipose stromal cells revealed increased macrophages displaying gene expression signatures of the alternately activated type. Mechanistically, depletion of such phagocytic immune cells in iAdFASNKO mice fully abrogated appearance of thermogenic adipocytes in iWAT. Altogether, these findings reveal an unexpected pathway of cAMP/PKA-dependent iWAT browning that is initiated by adipocyte signals and caused by macrophage-like cells independent of sympathetic neuron involvement.

## INTRODUCTION

It is well recognized that adipose tissue depots in rodents and humans can strongly influence systemic glucose and lipid homeostasis^1-3^. Thermogenic brown and beige adipocytes are especially active in this regard, as they can enhance energy expenditure as well as secrete potent factors that act on the metabolism of distant tissues^4-7^. Expansion of brown adipose tissue (BAT) and increased appearance of beige adipocytes in inguinal white adipose tissue (iWAT) of mice and humans during cold exposure is associated with remodeling of tissue architecture^8-10^, and is controlled by activation of local sympathetic nerve fiber (SNF) activity^11-14^. Single cell RNA transcriptomic analysis has corroborated the extensive cellular heterogeneity of adipose depots and identified various resident immune cells and other cell types that are present^15-20^. Moreover, the association between increased abundance of iWAT macrophages with anti-inflammatory, alternatively activated properties and cold-induced adipose remodeling has been demonstrated^15,21-23^. Norepinephrine (NE) released from SNFs activates the β-adrenergic receptor (βAR)-cAMP/PKA signaling pathway to induce these morphological and thermogenic changes during cold stimulus^24,25^. Accordingly, denervation of iWAT depots blocks cold-induced thermogenesis and appearance of beige adipocytes^26,27^. Overall, activation of this β-adrenergic pathway to modulate adipose tissue composition and functions yields increased glucose tolerance and resistance to high fat diet (HFD)-induced insulin resistance^24,28^.

Based on these beneficial metabolic effects of adipose browning, it is of interest to note that stimuli other than cold exposure can also mediate such effects^5,6^. These include intermittent fasting^29^, caloric restriction^30^, exercise^31^ and response to burns^32^. In addition, perturbations of metabolic pathways selectively within white adipocytes can trigger the appearance of beige adipocytes expressing uncoupling protein 1 (UCP1) in iWAT depots^33-36^. One such trigger of iWAT browning is the adipocyte-selective ablation of the last enzyme in de novo lipogenesis, fatty acid synthase (FASN), and this occurs even when the ablation is induced in fully mature mice^34-36^. Such selective ablation of adipocyte FASN in mice (iAdFASNKO) is accompanied by improved glucose tolerance and insulin sensitivity^34,36^. However, deletion of FASN in cultured adipocytes *in vitro* failed to cause UCP1 upregulation in the presence or absence of β-adrenergic stimulation^34^. Furthermore, data from this mouse model showed that signals emanating from FASN-deficient iWAT can affect distant BAT depots, presumably by transmission through the circulation or nervous system^35^. Similar to what occurs in cold induced iWAT browning, iAdFASNKO mice displayed increased expression of tyrosine hydroxylase (TH) in iWAT and BAT^34,35^ and increased sympathetic nerve activity in these tissues^35^. Taken together, these data suggested that in response to disruption of de novo lipogenesis adipocytes could be induced to release paracrine factors that might act locally to activate SNFs and initiate the appearance of beige adipocytes in iWAT.

The initial aim of the present studies was to determine whether SNFs within iWAT drive the emergence of beige adipocyte in iWAT by FASN-deficient adipocytes in mice, similar to cold exposure^11,12^. In marked contrast to the expected outcome, two independent denervation procedures that blocked cold induced iWAT browning failed to disrupt the appearance of beige adipocytes in iAdFASNKO mice. Instead, single cell transcriptomic analysis of the stromal vascular fraction of iWAT from iAdFASNKO mice revealed enhanced macrophage polarization towards the alternatively activated type known to be associated with tissue remodeling. The ablation of macrophages from iWAT of iAdFASNKO mice fully prevented the emergence of beige adipocytes. These data reveal the presence of an alternate pathway for adipocyte browning in iWAT that is impervious to denervation of adipose tissue but rather inhibited upon depletion of adipose tissue macrophages.

## RESULTS

### Imaging UCP1 upregulation in whole iWAT depots of iAdFASNKO mice

Based on our previous data indicating that, similar to cold exposure, induced deletion of FASN in adipocytes of mature mice (iAdFASNKO) enhances both beiging and innervation in iWAT^34,35^, we performed 3D imaging of entire iWAT depots (Fig. 1 and Extended Data Movies 1-5) to visualize the distribution of sympathetic nerve fibers (SNFs) and beige cells. Using the Adipo-Clear method for whole adipose clearing and 3D immunolabeling^11^, iWAT samples from wild type, iAdFASNKO (room temperature) and cold exposed wild type mouse were immunolabeled for uncoupling protein 1 (UCP1) and tyrosine hydroxylase (TH) and analyzed by lightsheet microscopy (Fig. 1a and Extended Data Movies 1-5). The 3D images show massive numbers of UCP1+ cells in iWAT from iAdFASNKO as well as the cold exposure group (Fig. 1 and Extended Data Movies 1-4). Strikingly, the UCP1+ cells were mostly abundant in the specific inguinal area of iWAT (Fig. 1a), similar to previously published data for cold exposed mice^11^. Interestingly, when imaged with 24x magnification, individual UCP1-positive multilocular adipocytes could easily be seen (Fig. 1b), and the abundance of such cells is somewhat less in the iAdFASNKO mice compared to cold exposed mice (Fig1a, b). Three-dimensional projections (Extended Data Movies 1-3) and optical sections with high power magnification (Extended Data Movies 4-5, 70μm and 10μm, respectively) of the iWAT depots also showed dense networks of SNFs detected by anti-TH in all groups. Although TH protein levels were increased in iWAT from iAdFASNKO mice when analyzed by western blot, we could not detect significant differences in the 3D projections of the sympathetic arborization when comparing control vs. iAdFASNKO or vs cold exposure using this method and limited samples.

**Figure 1.**
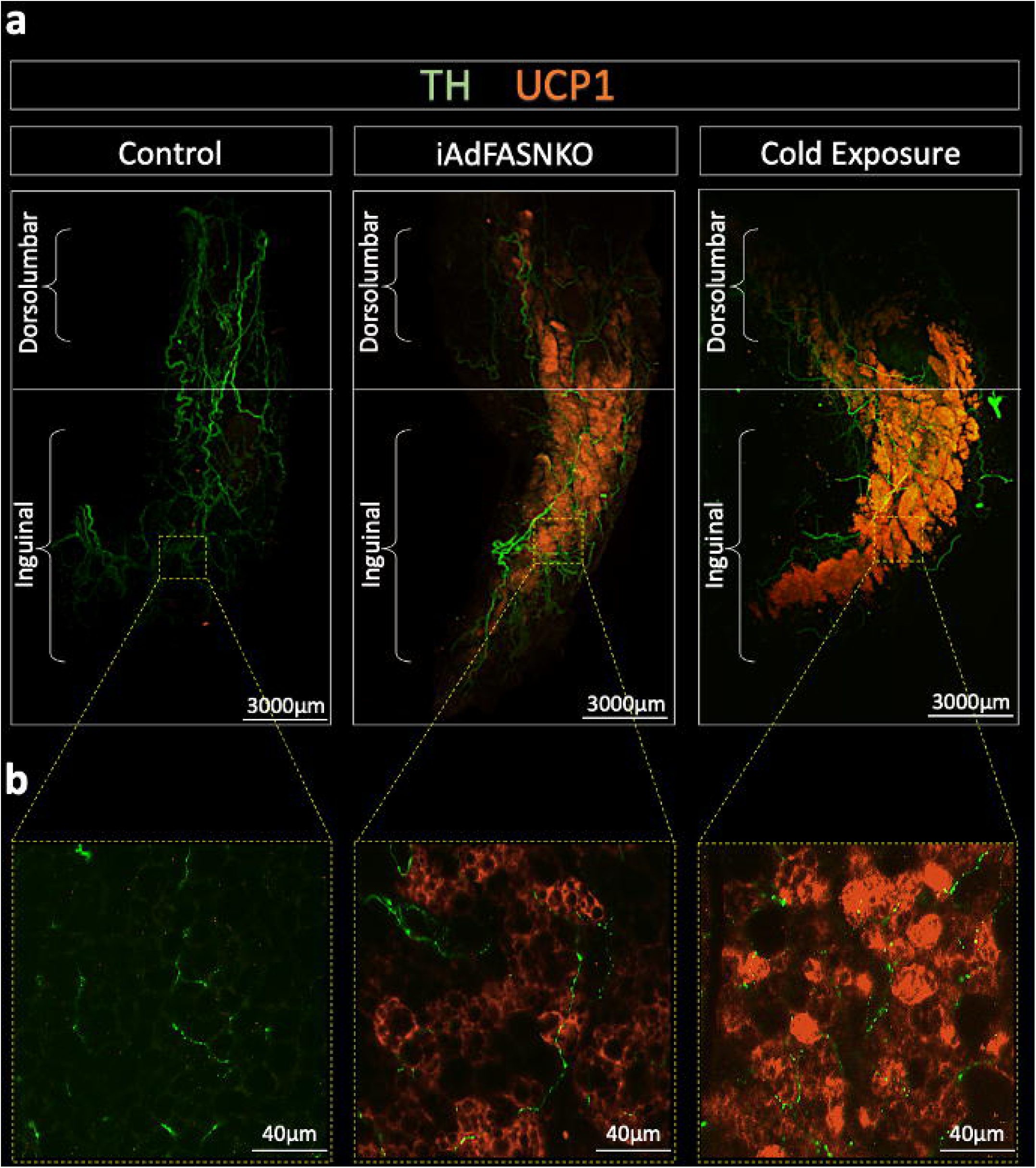
3D view of whole iWAT depots reveals robust UCP1 upregulation in the iAdFASNKO mouse. All panels are light-sheet fluorescence microscopy images of inguinal white adipose tissue (iWAT) processed using the Adipo-Clear technique. **(a)** Representative three-dimensional reconstruction of adipose sympathetic nerves and thermogenic profile of iWAT from control, iAdFASNKO, and 6°C cold-exposed mice. The samples are immunolabeled with anti-tyrosine hydroxylase (green) and anti-UCP1 (orange) and imaged at **(a)** 1.1× and **(b)** 24× magnification. Scale bars are provided for each panel.

### iWAT denervation does not block iAdFASNKO- induced adipose beiging

It is well known that cold-induced iWAT beiging in mice is absolutely dependent upon sympathetic innervation^11,12,37^. To determine whether sympathetic nerve fibers are essential for the browning of iWAT, we performed two procedures that disrupt local sympathetic innervation in iWAT, including a chemical approach using local iWAT 6-hydroxydopamine (6-OHDA) injection^37-39^ as well as local iWAT surgical denervation that it is known to destroy sympathetic and sensory nerves within the tissue^26,38,40^ (Fig. 2 and Extended Data Fig. 1b-e). As positive controls, sham and chemically denervated wild type mice were housed at 6°C for six days to induce iWAT beiging (Fig. 2c, d). Visual inspection of the iWAT depots in these mice show cold- and iAdFASNKO-induced shrinkage of the tissue and a strong browning effect (Fig. 2b, d). Strikingly and surprisingly, the iAdFASNKO-induced browning was not noticeably affected by iWAT denervation, while appearance of beige adipocytes in the cold-induced mice was blocked (Fig. 2b, d). The ability of chemical denervation to block cold-induced but not iAdFASNKO-induced UCP1 protein (Fig. 2e, g, h) and mRNA (Fig. 2f) confirmed these observations. Successful denervation in these experiments is shown by the decreased TH signal due to 6-OHDA treatment by Western blot analysis (Fig. 2g, h). Interestingly, the adipocytes in the cold-exposed mice display a smaller diameter when compared to the room temperature group (Fig. 2e). These results indicate that sympathetic innervation is essential for the cold-induced but not iAdFASNKO-induced appearance of UCP1-expressing beige adipocytes in iWAT.

**Figure 2.**
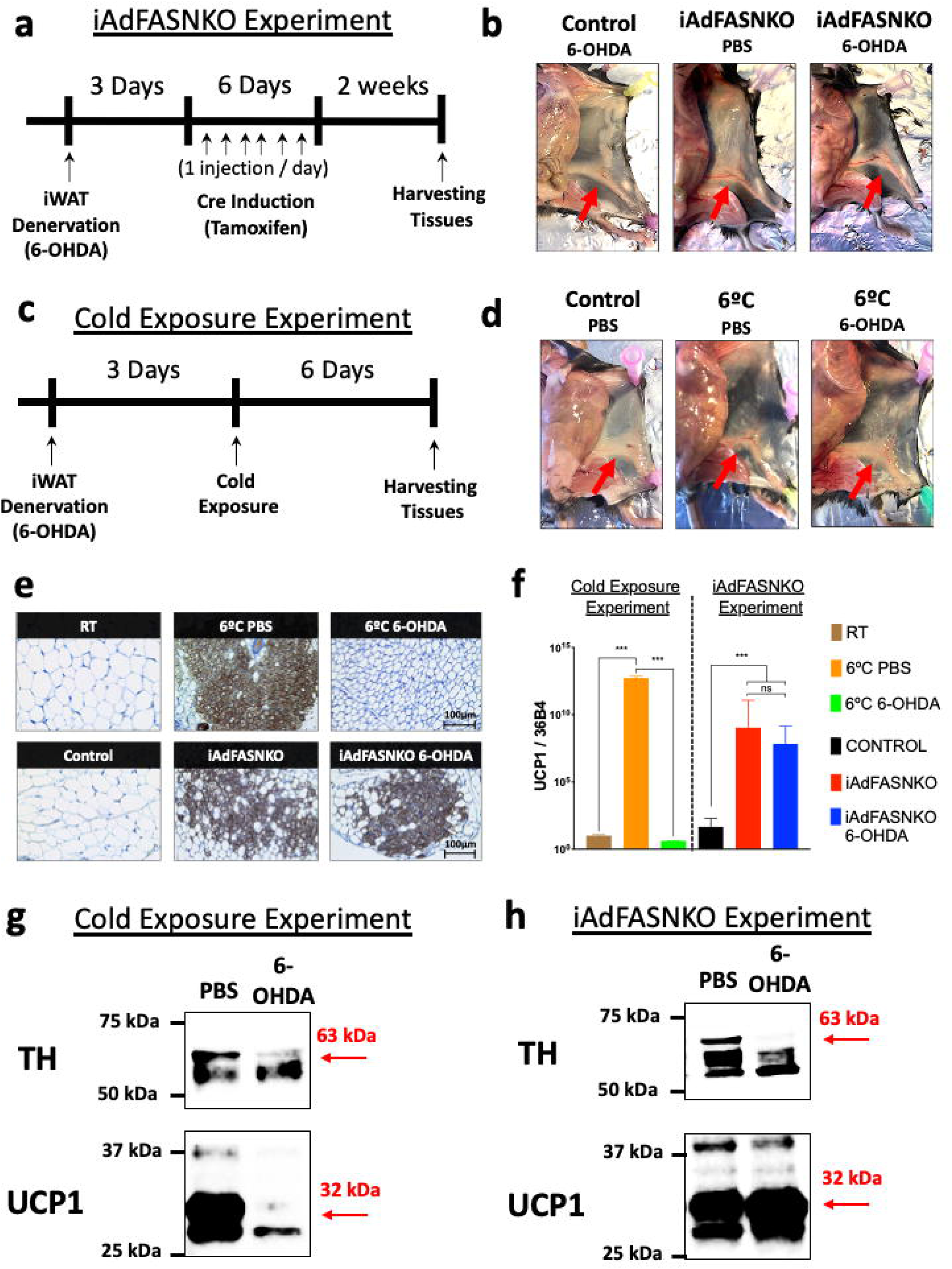
Denervation fails to block iWAT “browning” in response to iAdFASNKO. **(a,c):** Diagram representing the chemical denervation experiment in **(a)** iAdFASNKO mice and **(c)** cold-exposed wild-type mice. **(b,d):** Representative images of mice emphasizing the iWAT (see red arrows) after chemical denervation and **(b)** iAdFASNKO or **(d)** cold exposure. **(e):** Immunohistochemistry for detection of UCP1 in iWAT from cold-exposed and iAdFASNKO mice with and without chemical denervation. Scale bar is provided in the panel. **(f):** qRT-PCR was performed for *Ucp1* mRNA quantification in iWAT from same groups as **(e)**. N = 3- 4 per group. These results are representative of five independent experiments. The results are means ± SEM. *** P < 0.001, vs. control or room temperature (RT) group. **(g,h)** Depicted are representative westerns blot analyses for detection of TH and UCP1 protein levels in both experimental conditions (cold exposure **(g)** and iAdFASNKO **(h)**).

Another key finding in this series of experiments was that although chemical denervation blocked TH up-regulation, it did not blunt activation of the PKA pathway in response to iAdFASNKO (Extended Data Fig. 1a). Anti-phospho-PKA substrates, anti-phospho-HSL, and anti-phospho-perilipin staining remained elevated in iWAT from iAdFASNKO mice that were chemically denervated (Extended Data Fig. 1a). Therefore, 6-OHDA-treated iWAT not only did not interfere with UCP1 upregulation in the iAdFASNKO mice, but also failed to decrease the activation of the cAMP/protein kinase A pathway known to induce UCP1 expression^41,42^.

To confirm the unexpected results obtained with chemically denervated iWAT in iAdFASNKO mice, we employed a surgical denervation approach known to be effective at local tissues including iWAT^38,42,43^. As expected, surgically denervated iWAT failed to elicit cold-induced beige adipocytes in the context of a substantial reduction in TH protein levels (Extended Data Fig. 1b). Similarly, no UCP1 upregulation by cold exposure after surgical denervation of iWAT was found (Extended Data Fig. 1b, d), confirming that innervation is required for the cold-induced browning in iWAT. In contrast, and consistent with the results of chemical denervation, no decrease was observed in abundance of beige cells or UCP1 expression after surgical denervation in the iWAT from iAdFASNKO compared with sham iWAT (Extended Data Fig. 1c, e). Altogether, these data indicate that signals emanating from cell types other than sympathetic nerve fibers are directing the beiging of iWAT in iAdFASNKO mice.

### Nrg4 deficiency attenuates cold-induced but not iAdFASNKO-induced beiging

Previous work has shown uniquely high levels of Neuregulin 4 (Nrg4) secreted from brown and beige adipocytes compared to other cell types^44-47^, and it has been suggested that Nrg4 may enhance sympathetic nerve-induced beiging^47-49^. Based on this concept, coupled with the above data suggesting divergent pathways for promoting iWAT thermogenesis, we hypothesized that Nrg4 may be required for optimal iWAT beiging in response to cold exposure but not in iAdFASNKO mice. Indeed, when control versus Nrg4 whole-body knockout (NRG4KO) mice were exposed to 6°C temperature for six days, the beiging of iWAT was extensive in control mice but attenuated in NRG4KO mice (Fig. 3a-d). Thus, UCP1 mRNA expression (Fig. 3a) and protein expression (Fig. 3b-d) in iWAT were both decreased by about 50% in mice with Nrg4 deficiency. This was readily visualized in immunohistochemistry images of iWAT stained for UCP1 (Fig. 3b). In contrast, double KO mouse model, deficient in both Nrg4 and adipose tissue FASN displayed the same amount of iWAT beiging as iAdFASNKO mice (Fig 3 e-g). Both UCP1 mRNA and protein levels were highly and equally upregulated in the iAdFASNKO and the double KO mice (Fig 3 e-g). Thus, adipocyte deficiency of FASN causes optimal iWAT beiging independent of both iWAT SNFs and Nrg4, while optimal beiging of iWAT during cold exposure requires both sympathetic nerve activity and Nrg4.

**Figure 3.**
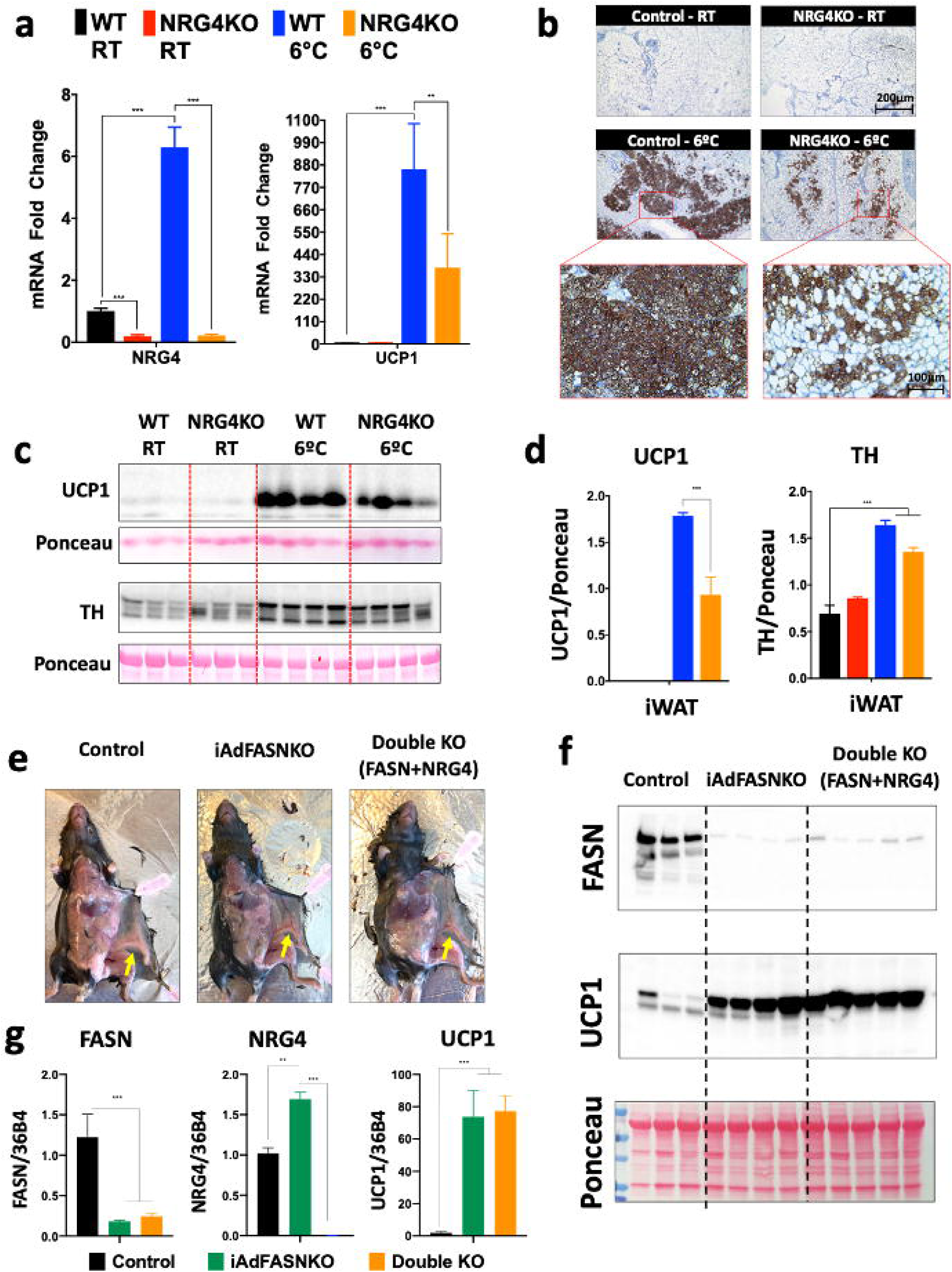
NRG4 deficiency in mice attenuates cold-induced but not adipocyte FASNKO-induced iWAT browning. **(a):** qRT-PCR was performed for *Nrg4* and *Ucp1* mRNA quantification in iWAT from control and NRG4KO mice housed either at 22°C or 6°C. N = 3-4 per group. The data are presented as the mean ± SEM. *** P < 0.001 and ** P < 0.05. **(b):** Depicted are immunohistochemical analyses for detection of UCP1 protein in iWAT in the different groups. **(c)** Western blots of iWAT lysates from control and NRG4KO mice housed either at 22 °C or 6°C. Shown are UCP1, TH and Ponceau staining (as a loading control) and **(d)** their respective protein quantifications. N = 3-4 per group. The data are presented as the mean ± SEM. *** P < 0.001. **(e):** Representative images emphasizing the morphology of the iWAT (see yellow arrows) of Control, iAdFASNKO and Double KO (FASN+NRG4) mice. **(f)** Western blotting for the detection of FASN and UCP1 protein levels in iWAT for the indicated mouse line. **(g):** qRT-PCR was performed for *Fasn, Nrg4* and *Ucp1* mRNA in iWAT from control, iAdFASNKO and Double KO (FASN+NRG4) mice. N = 5-8 per group. The results are means ± SEM. *** P < 0.001 and ** P < 0.05 vs. control group.

### Single-Cell RNA-Seq reveals increased iWAT alternatively activated macrophage content in iAdFASNKO mice

Since sympathetic innervation is not required for adipose beiging in iAdFASNKO mice (Fig. 2) and cell autonomous effects also appear not to play a major role^34^, we hypothesized that other adipose resident cell types may be targets of FASN-deficient adipocyte signals. We therefore applied single cell RNA-seq, a powerful technique to identify cell types and their mRNA expression profiles within complex tissues including adipose tissues from cold exposed mice^15,19,20,50^, to this question. The stromal vascular fractions (SVF) of iWAT from control and iAdFASNKO mice were isolated and subjected to RNA-seq analysis (see single cell RNA-seq workflow for more details related to the protocol in Extended Data Fig. 2a). To better resolve the relatively low numbers of immune cells present in the SVF in control mice, SVF cells were isolated and fractionated into lineage marker-positive (Lin+), mostly immune cells, and all other stromal cells (Lin-) using magnetic bead cell sorting (MACS) as reported by the Granneman laboratory^15^. Thus, 4 different libraries were prepared from an estimated approximately 6,000 cells per condition (Control and iAdFASNKO, Lin+ and Lin-), and were subjected to sequencing using Illumina NovaSeq 6000 to obtain 2.5 billion reads per run, yielding ∼625 million reads per sample. This number of reads per sample (∼100,000 reads per cell) was important to gain greater depth of coverage for detecting rare cell populations.

In analyzing the sequencing results from the above experiment, we first aggregated (Lin+ and Lin- libraries from control and iAdFASNKO) and normalized the data, and subjected it to graph-based clustering to identify cell types/states, which were projected onto the UMAP plot using the Loupe Cell Browser (Fig. 4a). Single cell RNA-seq data from the aggregated libraries enabled us to distinguish eight distinct clusters (Fig. 4a, b). Using differential gene expression between clusters, we analyzed the top 20 upregulated genes that we can identify and named each different cell cluster. The Violin plots in Extended Data Fig. 3 show the representative top upregulated genes from the different clusters, further revealing that each cluster also uniquely expresses specific marker genes. The total percentage of the eight clusters from the aggregated data set that were identified and the representative top up-regulated genes are: Collagen-Rich Progenitors, 51.8% (Col4a2), Pi16+ Progenitors, 19.2% (Pi16), Endothelial Cells, 4.5% (Vwf), Schwann Cells, 3% (Mpz) Smooth Muscle Cells, 2.9% (Acta2), Macrophage-Like, 5.13% (Lyz2), T cells, 6.9% (Ccl5) and B Cells, 6.2% (Cd79a) as illustrated in Fig. 4b. Segregation of the aggregated UMAP plot revealing differences among the number of cells in the different clusters demonstrated that the adipose tissue environment present in SFV from iAdFASNKO mice appears to have a somewhat different composition compared to the control group (Extended Data Fig. 2b).

**Figure 4.**
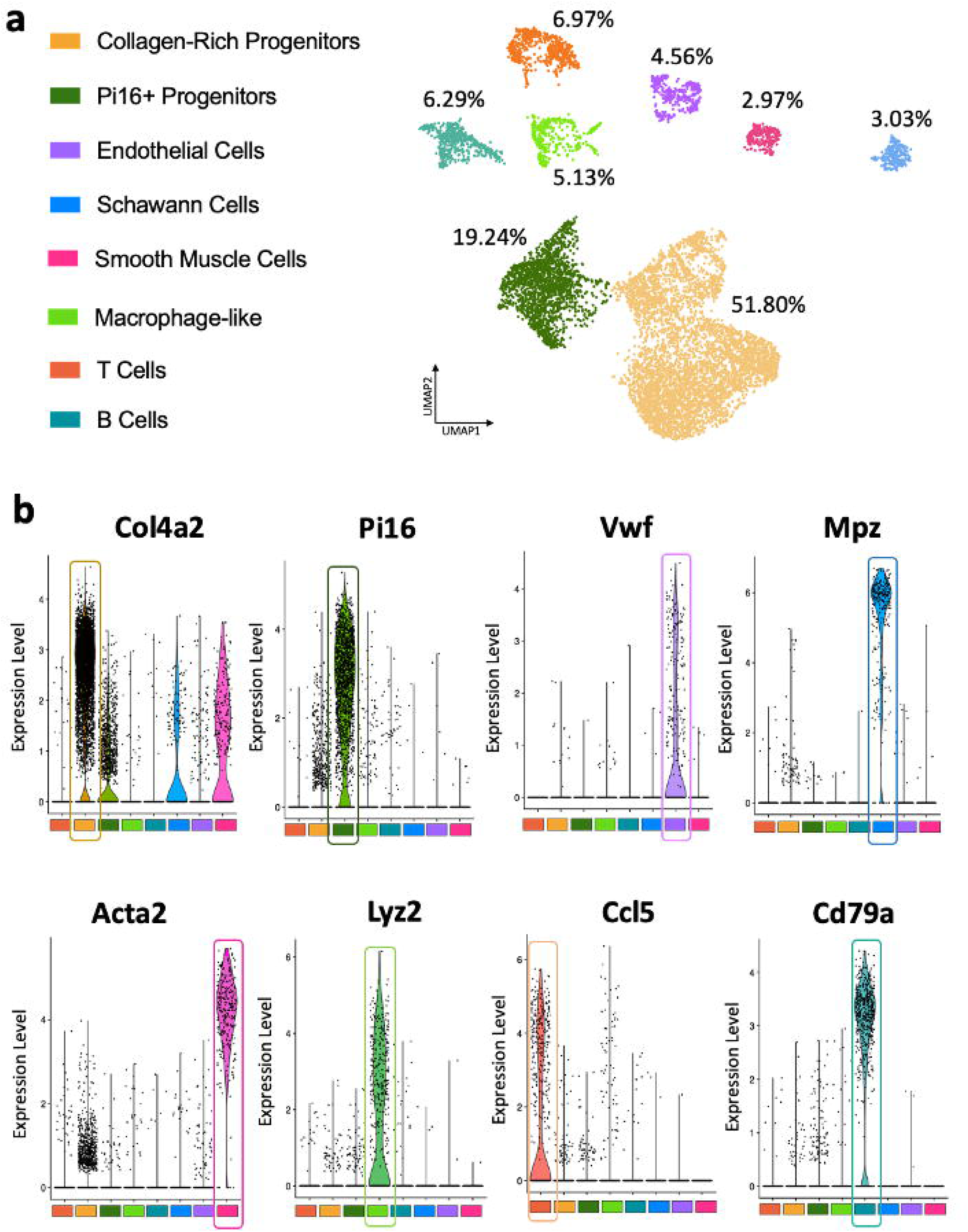
Single-Cell RNA-Seq reveals heterogeneity of non-adipocyte cell types within iWAT from wild type and iAdFASNKO mice. **(a)** Aggregated UMAP plot (control lineage-positive, control lineage-negative, iAdFASNKO lineage-positive and iAdFASNKO lineage-negative) of 18,336 stromal vascular cells (∼5,000 cell per condition) from iWAT showing eight different clusters (color code). Clusters were generated in the 10× Cell Ranger software via K-means clustering based on the PCA of the transcriptomic signature. Cluster names were manually determined based on the top-20 upregulated genes related to differential gene expression between clusters and knowledge of canonical cell markers. **(b)** Violin plots of log_e_ (counts/cell normalized to 10,000) for the top gene from each cluster in the aggregated dataset. Each point represents the log-expression value in a single cell. These representative genes were used to identify and name the following clusters: Collagen-Rich Progenitors (Col4a2), Pi16+ Progenitors (Pi16), Endothelial Cells (Vwf), Schwann Cells (Mpz), Smooth Muscle Cells (Acta2), Macrophage-like (Lyz2), T cells (Ccl5) and B Cells (Cd79a).

Based on numerous publications^21,51-55^ that have brought attention to the concept that macrophage may play a major role in adipose tissue remodeling and thermogenesis, we subjected the Macrophage-like cluster to a sub-clustering process, using only Lin+ libraries from control and iAdFASNKO mice to maximize the detection of rare population types (Fig. 5). Interestingly, this sub-clustering revealed five new sub-clusters related to the original cluster (Macrophage-Like) that were not displayed before using the aggregate UMAP plot. Applying the same approach described above to identify and name these new sub-clusters, we uncovered the following clusters: Resident Macrophage, M1 Macrophage, M2 Macrophage, Neutrophils cells and Dendritic Cells (Fig. 5a and Extended Data Fig. 4). Heatmaps of graph-based log2 fold change shows the top 5 up-regulated differentially expressed genes between sub-clusters (Fig. 5a). Segregation of the sub-cluster UMAP plot shows marked changes in the number of cells in the different clusters when comparing control vs. iAdFASNKO mice (Fig. 5b).

**Figure 5.**
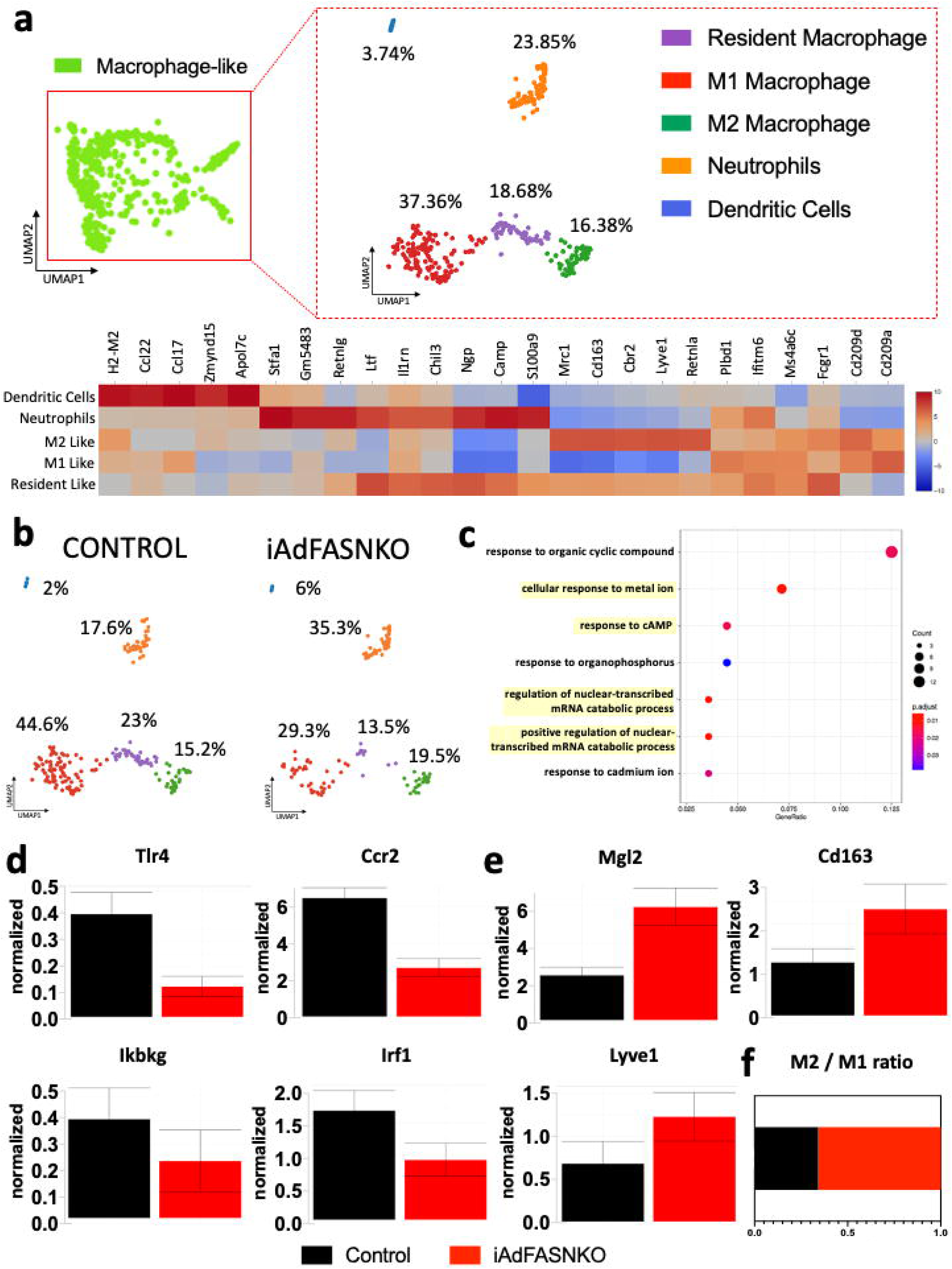
iWAT from iAdFASNKO reveals a shift in the M1- to M2- macrophage polarization in the macrophage-like cluster. The Sub-clustering dashboard of Macrophage-like cells from the aggregated clusters. **(a)** Lineage positive cells in the Macrophage-like cluster from figure 4 were clustered into five new sub-clusters via visual inspection within the Loupe viewer. The heatmap shows the top 5 differentially expressed genes between sub-clusters obtained from graph-based log2 fold changes generated from the Loupe browser. Colors (values) are z-score of the average log fold-change between sub-clusters for each gene (so z I is based on fold change across the 5 sub-clusters). Cluster names were manually determined based on the top-10 upregulated genes related to differential gene expression between clusters and knowledge of canonical cell markers. **(b)** UMAP-plot from (a) split into the different conditions: Control Lineage positive vs. iAdFASNKO Lineage positive. **(c)** Gene ontology (GO) and KEGG pathway enrichment (biological processes, the adjusted p-value of < 0.05) among upregulated differentially expressed marker genes in iAdFASNKO related to M2 macrophage cluster (Fig. 4c). **(d)** log_e_ of normalized expression values for M1- (Ccr2, Tlr4, Ikbkg and Irf1) and **(e)** M2- (Mgl2, Cd163 and Lyve1) macrophage polarization markers. Graphs show the mean ± SEM. **(f)** The ratio of M2/M1 adipose tissue macrophage in the stromal vascular cells from iWAT from control vs. iAdFASNKO.

To examine the possible function of each unique subpopulation, we investigated gene ontology (GO) and KEGG pathway enrichment of the upregulated differentially expressed marker genes in iAdFASNKO related to the M2 macrophage cluster (Fig. 5c). This analysis shows that the M2 macrophage population in iAdFASNKO iWAT SVF has an enrichment in genes related to pathways that could be linked to regulation of nuclear-transcribed mRNA catabolic processes and responses to cAMP (Fig. 5c). Furthermore, iAdFASNKO elicits an intense adipose tissue macrophage polarization toward the M2-state, demonstrated by downregulation in general markers of inflammation (Tlr4, Ccr2, Ikbkg and Irf1 - Fig. 5d) and a robust up-regulation in M2 macrophage markers (Mgl2, Cd163 and Lyve1 - Fig. 5e). This leads to a higher ratio of M2/M1 macrophages in the SVF from iAdFASNKO (Fig. 5f), implying that M2 macrophage polarization may be involved in the molecular mechanism of how iAdFASNKO enhances thermogenesis in iWAT. This concept is reinforced by studies suggesting that alternatively activated macrophages (M2 macrophage polarization) are involved in the iWAT beiging process and play an essential role in maintaining systemic metabolism^14,22,30,54,56,57^. All the significant (p<0.05) up- regulated and down-regulated genes in iAdFASNKO vs control related to the Macrophage-like sub-clustering are present in the Source Data 1 related to the Fig. 5.

### Adipose iAdFASNKO enhances macrophage polarization towards the alternatively activated type in iWAT

In further characterizing the macrophages in iWAT, we performed histological analyses that showed a positive signal for F4/80 immunostaining among the iWAT multilocular adipocytes in the iAdFASNKO and iAdFASNKO+ denervation groups (Fig. 6a). Similarly, we found F4/80 positive cells among the beige adipocyte areas in the iWAT after six days of cold exposure (Extended Data Fig. 5d) in wild type mice, comparable to what has been reported found by other groups^21,58^. The specificity of the F4/80 antibody was confirmed by the detection of adipose tissue macrophage infiltration in the iWAT from ob/ob mice (Extended Data Fig. 5e).

**Figure 6.**
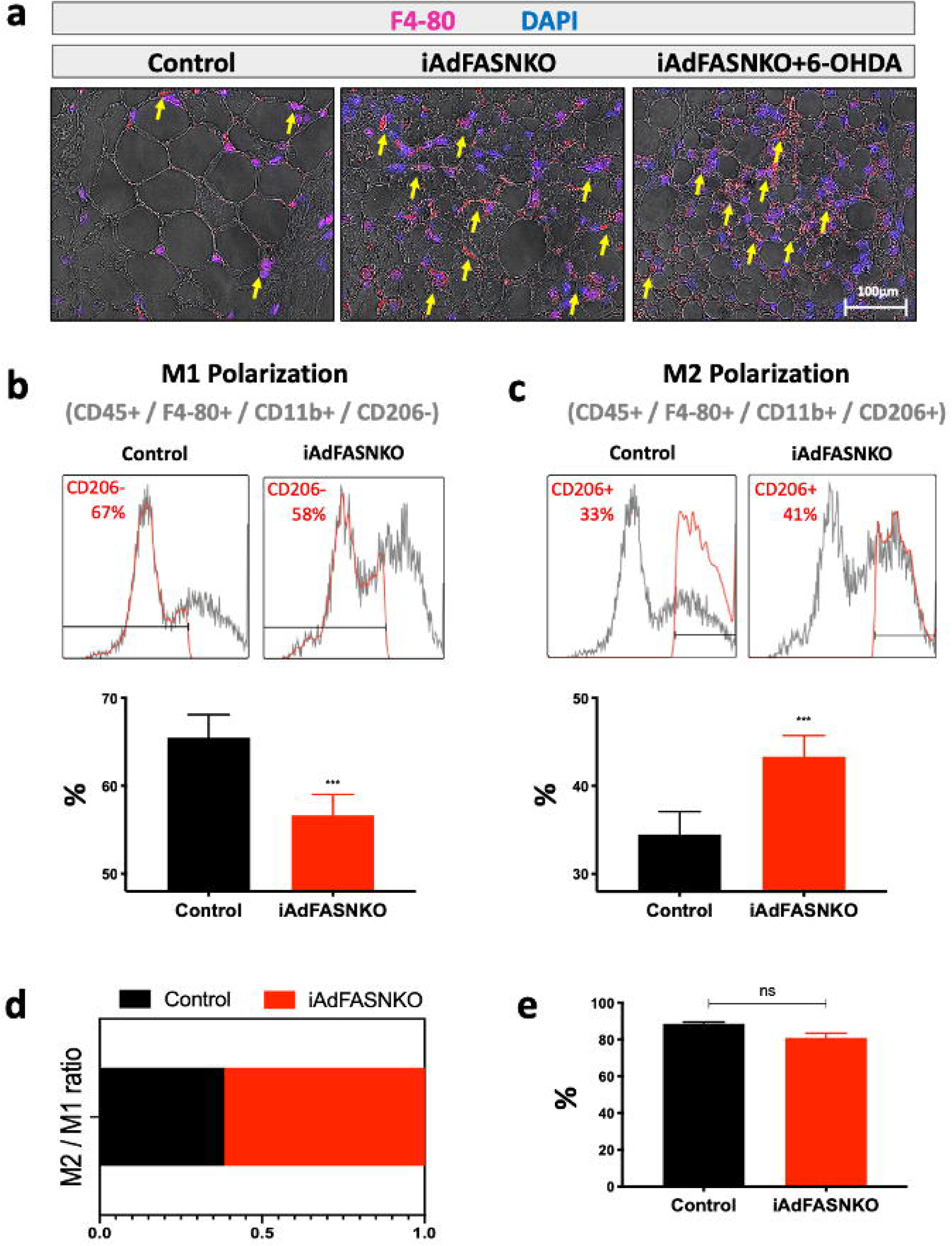
FACS analysis confirms the enrichment of M2-type macrophage populations in the iWAT of iAdFASNKO mice. **(a):** Representative images for immunofluorescent staining of paraffin-embedded tissue sections for pan- macrophage marker F4/80 (red), DAPI (blue) and bright-field (BF, grey) in the iWAT from the different experimental groups (Control, iAdFASNKO and iAdFASNKO+6-OHDA). Yellow arrows indicate cell F4/80+ cells located in the browning area (multilocular adipocytes) of iWAT from the iAdFASNKO and iAdFASNKO+6-OHDA group. **(b-c):** Stromal vascular fractions (SVF) were isolated from iWAT by collagenase digestion for each different group. Flow cytometric analysis of SVF was conducted using fluorescent-conjugated antibodies against CD45, F4/80, CD11b and CD206. Adipose tissue macrophages were defined as CD45^+^F4/80^+^CD11b^+^. **(b)** M1 and **(c)** M2 macrophages were defined as CD45^+^F4/80^+^CD11b^+^CD206^-^ and CD45^+^F4/80^+^CD11b^+^CD206^+^, respectively. **(d)** M2/M1 adipose tissue macrophage ratio in the iWAT from control vs. iAdFASNKO. **(e)** The total number of macrophages was defined as CD45^+^F4/80^+^CD11b^+^. Representative results of flow cytometry are shown. N = 4 per group. Graphs show the mean ± SEM. Two-way ANOVA determined statistical significance, ***P < 0.001.

To validate the data generated by our single cell RNA-seq analyses, flow cytometry methods were performed on the SVF from iWAT to evaluate the adipose tissue macrophage polarization state in iAdFASNKO mice. In these analyses, CD45+/F4/80+/CD11b+/CD206− cells were marked as M1-positive cells, and CD45+/F4/80+/CD11b+/CD206+ cells as M2-positive cells (Fig. 6b, c). These FACS studies provided data corroborating that obtained from single cell RNA-seq, as the SVF from iAdFASNKO iWAT exhibited a reduction in M1 macrophage content, and an enhancement in the M2 macrophage population (Fig. 6b, c). The resulting increased ratio of M2/M1 macrophages in the iWAT from iAdFASNKO (Fig. 6d) was statistically significant and similar to what was observed from the single cell RNA-seq data.

In contrast to the large difference in the M2/M1 iWAT macrophage ratio between control and iAdFASNKO mice, there was no difference in the total number of macrophage cells detected between these different groups of mice (Fig. 6e). Similarly, no differences were observed in apparent cell proliferation rates in the SVF between control and iAdFASNKO mice, as detected by labeling the SVF cells with 5-ethynyl-2′-deoxyuridine (EdU) *in vivo*. Thus, flow cytometry analyses of EdU incorporation failed to show any difference in proliferation ratio in any of the cell types analyzed (Immune cells - EdU+/CD45+, Progenitors cells - EdU+/PDGFRα and Endothelial cells - EdU+/CD31+) between iAdFASNKO mice and the control group (Extended Data Fig. 5a-c). Taken together, our analyses indicate a strong association between remodeling in the iWAT and M2 macrophage polarization in iAdFASNKO mice.

### iWAT macrophage depletion impairs iAdFASNKO-induced browning

To test the hypothesis that iWAT M2 macrophage enhancement in iAdFASNKO mice plays a physiological role in the induction and maintenance of iWAT thermogenesis, we administered clodronate-containing liposomes to deplete adipose tissue macrophages in the iWAT of iAdFASNKO mice (Fig. 7a). Clodronate liposomes were previously shown to selectively target and deplete phagocytic cells of the mononuclear phagocyte system (effectively monocytes/macrophages)^59-62^. This approach for adipose tissue macrophage depletion has been used successfully before by others^63-65^. As expected, endogenous Fasn mRNA was efficiently decreased in iWAT of iAdFASNKO and iAdFASNKO + clodronate mice (Fig. 7b). Gene expression analysis for macrophage markers was performed to validate the macrophage depletion in the iWAT. Clodronate liposomes effectively blunted the expression of F4/80, Cd206, Cd11c, and Cd68 only in the iAdFASNKO + clodronate group, confirming that iWAT macrophage depletion was successful (Fig. 7c). Importantly, iWAT macrophage depletion completely blocked the upregulation of UCP1 mRNA (Fig. 7d) and protein levels (Fig. 7e) in the iAdFASNKO group. Furthermore, multilocular adipocytes were not detected in iWAT of the iAdFASNKO + clodronate mice. Taken together, these findings are consistent with the concept that M2 macrophage polarization may be required to promote browning of iWAT in the iAdFASNKO mouse model.

**Figure 7.**
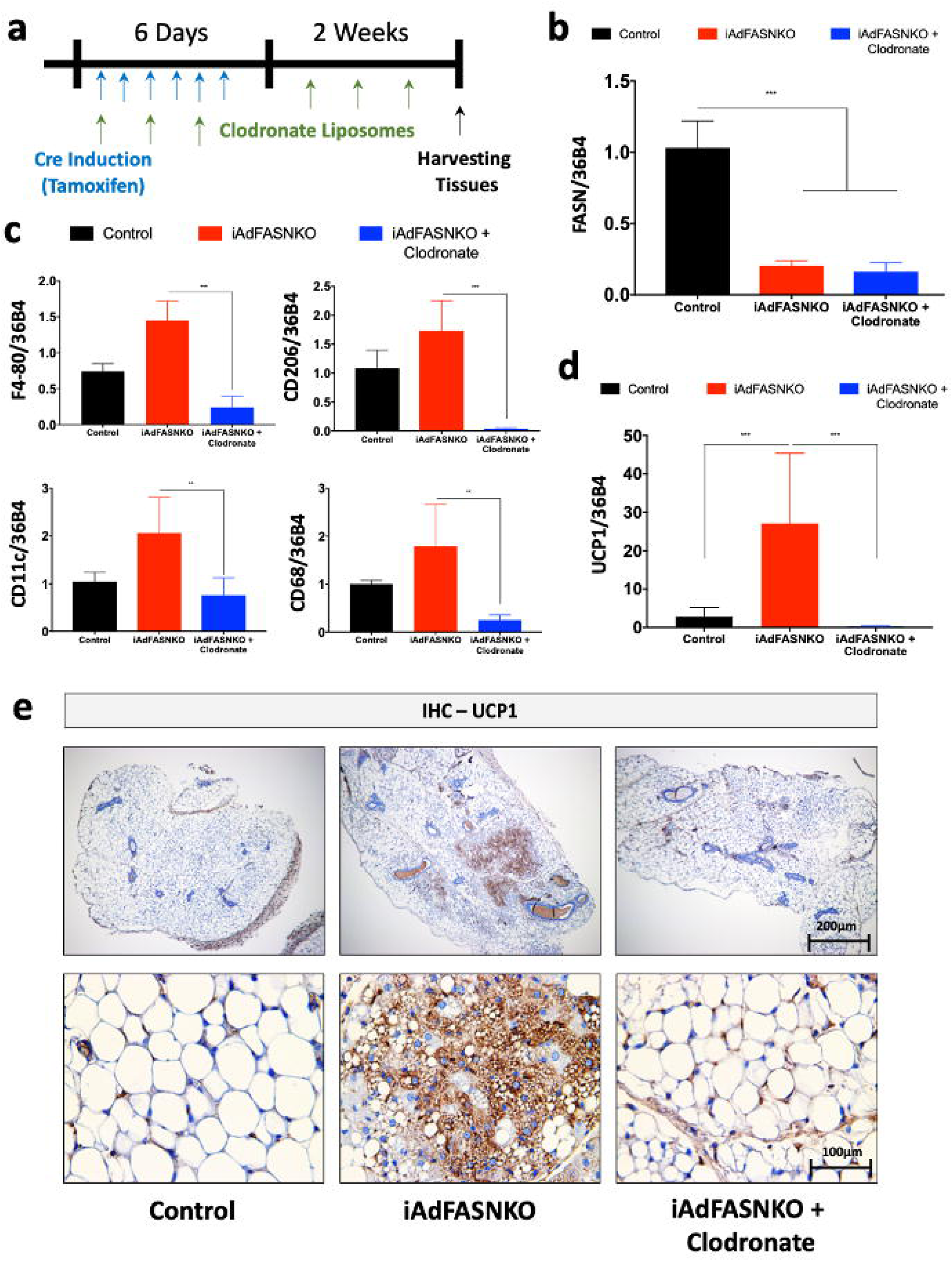
Depletion of macrophages blocks iAdFASNKO-induced browning. **(a):** Schematic representation of the experimental design for phagocytic immune cells depletion in the iAdFASNKO model using clodronate-containing liposomes. Every other day, mice received intraperitoneal injections of liposomes containing PBS or clodronate during a two-week period. **(b-d):** Shown are the qRT-PCR analyses to quantify mRNA expression levels of *Fasn, F4/80, Cd206, Cd11c, Cd68* and *Ucp1* in iWAT from control, iAdFASNKO and iAdFASNKO + clodronate. N = 6 mice per group. The data are presented as the mean ± SEM. ** P < 0.01; *** P < 0.001. **(e):** Immunohistochemistry was used for the detection of UCP1 protein levels in iWAT from control, iAdFASNKO and iAdFASNKO + clodronate. UCP1 staining reveals suppression of browning in iAdFASNKO after macrophage depletion. Representative images are shown. Scale bars are provided in the panel.

### Adipocyte cAMP signaling is required for iAdFASNKO-induced browning in iWAT

The cAMP/PKA signaling pathway is necessary for cold-induced browning of iWAT^25,66^, but the independence of adipose browning in iAdFASNKO mice from SNF innervation raised the question whether the cAMP pathway is involved. Stimulatory G protein (Gsα) activation is required for receptor activated increases in the production of cAMP by adenylyl cyclase, leading to activation of PKA, the phosphorylation of proteins and functional responses^24,28^. In order to determine if cAMP signaling is required for iAdFASNKO-induced browning, we generated a new animal model by crossing mice with genetic deletion of Gsα selectively in adipocytes (Adipo-GsαKO)^25^ with iAdFASNKO mice. Confirming the double KO, FASN and Gsα protein levels were shown to be reduced in iWAT of these double KO mice (Fig 8b). Loss of cAMP production by adipocyte inactivation of Gsα completely blocked the stimulatory effect of iAdFASNKO on Ucp1 and Cidea mRNA expression levels in IWAT (Fig. 8c). In agreement with the Ucp1 mRNA levels in Fig. 8c, the formation of UCP1-positive multilocular adipocytes was abolished in iWAT from Double KO (FASN+Gsα), as illustrated in Fig. 8d. Thus, these results strongly indicate that cAMP signaling in adipocytes is necessary for iAdFASNKO-induced browning in iWAT.

**Figure 8.**
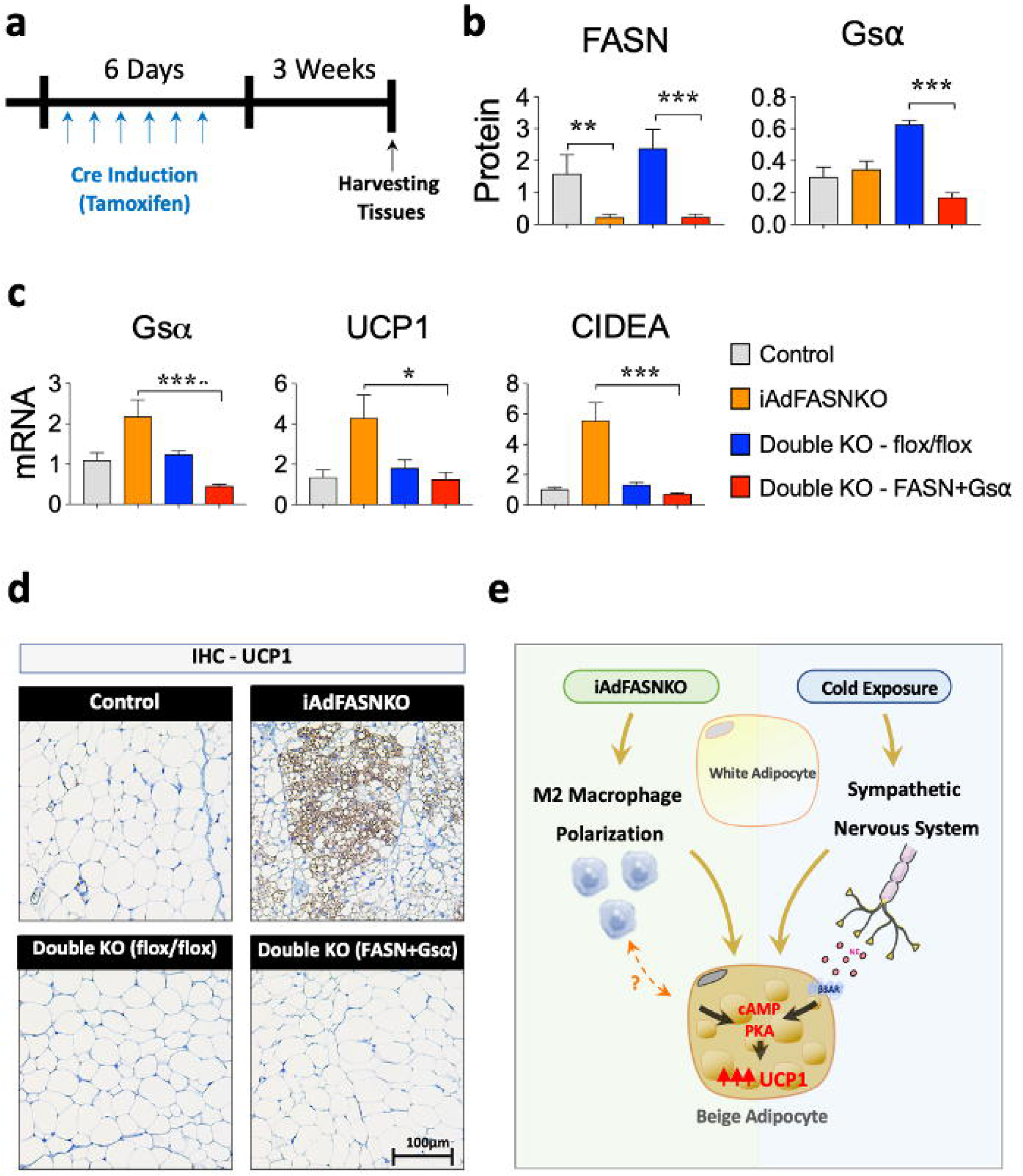
Adipocyte cAMP Signaling is Required for iAdFASNKO-induced browning in iWAT. **(a):** Diagram representing the Double KO (FASN+Gsα) experimental protocol. **(b):** FASN, Gsα and UCP1 protein levels of iWAT were quantified by densitometry from the immunoblot data from the different experimental conditions (Control, iAdFASNKO, Double KO (flox/flox) and Double KO (FASN+Gsα). GAPDH was used for loading control. **(c):** qRT-PCR was performed for *Gs*α, *Ucp1* and *Cidea* mRNA quantification in iWAT from different experimental conditions. N = 4-5 per group. The results are means ± SEM. *** P < 0.001, iAdFASNKO vs. Double KO (FASN+Gsα). ** P < 0.001, Control vs iAdFASNKO. * P < 0.05, iAdFASNKO vs. Double KO (FASN+Gsα). **(d):** Representative IHC analyses for the detection of UCP1 in iWAT from the different experimental conditions. Representative images are shown. Scale bars are shown in the panel. **(e):** The proposed model shows a divergence in signaling pathways sufficient to cause adipose beiging in response to cold exposure versus iAdFASNKO. In the latter case, the data suggest that adipose beiging signaling is SNS-independent and relies on the crosstalk between adipocytes and M2 macrophages in iWAT.

## DISCUSSION

The major finding of this study is that at least two divergent pathways involving adipose stromal vascular cell types are fully and independently able to cause appearance of thermogenic beige adipocytes within iWAT depots in mice. One pathway can be triggered by cold exposure and is dependent upon active sympathetic innervation of iWAT depots (Figs. 1 and 2), while a separate pathway initiated by adipocyte FASN deficiency is dependent upon phagocytic cells within iWAT but independent of innervation (Fig. 2). Interestingly, stimulation of iWAT beiging by both pathways are associated with increased sympathetic activity in iWAT^11,12,34,67^, even though it is not required for beiging in response to adipocyte FASN deficiency. It may be that the sympathetic activation in iAdFASNKO mice also stimulates appearance of beige adipocytes in iWAT, but that this pathway is redundant with the macrophage dependent pathway and not additive. Our imaging of iWAT by light sheet microscopy shows similar high-density sympathetic nerve fibers within the areas of beige adipocytes, both in cold-exposed and iAdFASNKO mice, but direct comparison of the sympathetic nerve activities in these two models has not been determined. The extent of the increase in iWAT sympathetic nerve density by cold exposure is controversial, as one group reports large increases^12^ while another reports little effect^11^. Our data from measuring total TH expression measured in entire iWAT pads suggests only modest increases due to either cold exposure (Fig. 3c, d) or adipocytes deficient in FASN^35^, in the range of two fold, which may not be easily quantified in the imaging experiments. Since the extent of beiging in iWAT is greater in cold exposure than in iAdFASNKO mice (Figs. 1 and 2), it is likely that cold exposure stimulates sympathetic nerve fiber activity more strongly than occurs in iAdFASNKO mice.

The involvement of adipose tissue macrophages in iWAT browning has been controversial following published evidence supporting the novel concept that macrophages themselves secrete catecholamine to activate adipocytes towards thermogenic activity^68,69^. Strong evidence against this hypothesis was then raised^70^, and more recently two groups reported that a specific macrophage population associated with sympathetic nerve fibers instead takes up norepinephrine within a degradation pathway^53,55^. Importantly, all of the reports that have implicated macrophage involvement in adipose tissue browning have done so in the context of intact sympathetic neurons, and no previous data that we know of has shown macrophages can mediate browning without such adipose tissue innervation^53-55^. Thus, the results presented here represent an advance on the issue of macrophage involvement in adipose browning, as we show it can operate effectively independent of sympathetic activation (Fig. 2).

Single cell transcriptomic analysis has been a powerful tool in identifying and quantifying the various cell types within adipose tissues^15-20^, and we therefore employed this method to probe the key question of what adipose cell types might be involved in mediating the phenotype of iAdFASNKO mice. By obtaining approximately 10^5^ sequencing reads per cell from adipose tissue SVF in our analysis, we were able to initially define adipose tissue cell types reasonably compatible with previous reports^15,19,20^, including endothelial cells, several immune cell types, Schwann cells, smooth muscle cells and progenitor cells. Interestingly, we noted an apparent increase in the abundance of a cell cluster we denote as peripheral neuron-associated Schwann cells based on their gene expression signature (Fig. 4 and Extended Data Fig. 3), similar to the increase we have reported for SNFs^34,35^. However, since SNFs are not required for adipose browning in iAdFASNKO mice, we focused our attention on the other cell type that has been repeatedly reported to change in cold-induced browning-macrophages^21,51-53^. Specifically, adipose browning is reported to be associated with an increased macrophage polarization towards what has been denoted as the alternatively activated M2 state^15,21-23^, and we observed a similar apparent increase in the M2 macrophage sub-cluster in iAdFASNKO mice(Fig. 5). This finding was confirmed independently by FACS analysis in iAdFASNKO mice (Fig. 6). These data combined with the requirement for iWAT macrophages to elicit adipose browning in iAdFASNKO mice (Fig. 7) suggests these cells are critical for mediating this iWAT browning.

Interestingly, the divergent pathways that mediate iWAT beiging, emanating from nerve fibers in cold exposed mice vs macrophages in iAdFASNKO mice, appear to converge at the level of the cAMP/PKA signaling pathway within the iWAT adipocytes. Thus, greatly increased abundance of phospho-perilipin and phospho-hormone sensitive lipase, known substrates of PKA^71,72^, are observed in the iWAT of iAdFASNKO mice compared to wild type mice (Extended Data Fig. 1a), similar to what is observed in cold exposed mice^73,74^. In the case of cold exposure, it is known that catecholamines emanating from sympathetic nerve fibers act through β-adrenergic receptors to stimulate the cAMP/PKA pathway to activate protein kinase p38 and downstream transcription factors that regulate expression of UCP1 and other thermogenic genes^28^. In contrast, in the case of iAdFASNKO mice, the data presented here suggest that catecholamines derived from local iWAT nerve fibers are not the cause of cAMP/PKA stimulation since it is not blocked by denervation (Extended Data Fig. 1a). However, the important experiment in Fig. 8 does show that the ability to generate cAMP within adipocytes is necessary to attain iWAT beiging in iAdFASNKO mice, as adipocyte-selective Gsα deficiency attenuates the expression of iWAT UCP1 in iAdFASNKO mice, as in cold exposure^25^. These data also reinforce the concept that beiging in iAdFASNKO mice results from direct conversion of white adipocytes to beige adipocytes since it is the cAMP in mature adipocytes that is necessary for the beiging (Fig. 8). Taken together, the data indicate that divergent inputs into cAMP generation in adipocytes lead to the similar outcome of iWAT beiging in the two models studied here.

It is not known what factors derived from alternatively activated macrophages might be most important in driving the emergence of beige adipocytes in iWAT from either cold exposure or in iAdFASNKO mice. One reported factor may be acetylcholine from immune cells, but this is not apparently sufficient to maintain chronic iWAT browning and appears to be derived from lymphocytes^52^. Suggestions that macrophages themselves can synthesize and secrete catecholamines to produce adipose browning^68,69^ have been dismissed by others^70^, but this idea remains controversial with conflicting data^22,75,76^. Clearly, a key gap in our knowledge about the connection between adipose tissue immune cells and emergence of iWAT beige adipocytes is how these cells mediate such effects, an important area of current investigation. However, an important conclusion from our present studies is that strong adipose browning and associated improvements in glucose tolerance can be elicited through mechanisms initiated by adipocytes themselves that do not require adipose tissue innervation by SNFs.

## METHODS

### Animal studies

Four-week-old male C57BL/6J (WT) mice were obtained from Jackson Laboratory. Mice were housed on a 12h light/dark schedule and had free access to water and food. Mice with conditional *Fasn*^flox/flox^ alleles were generated as described elsewhere^36^. To knockout *Fasn* in adult mice, homozygous *Fasn*^flox/flox^ animals were crossed to Adiponectin-Cre-ERT2 mice as previously described^34^. Briefly, at eight weeks of age, both *Fasn*^flox/flox^ and *Fasn*^flox/flox^ -Adiponectin-Cre-ERT2 (iAdFASNKO) were treated once a day with 1 mg tamoxifen (TAM) dissolved in corn oil via intraperitoneal (i.p.) injection for six consecutive days. *Nrg4* whole-body knockout (NRG4KO) mice were provided by Dr. Jiandie D. Lin. To obtain the double knockout mouse model, iAdFASNKO was bred with NRG4KO to generate an iAdFASNKO + NRG4KO mouse line. Adipocyte deletion of FASN in iAdFASNKO + NRG4KO was achieved by treating adult mice with TAM injections as described. To selectively delete FASN and Gsα in adipocytes from adult mice, TAM-inducible adiponectin-GsαKO were crossed to iAdFASNKO mice to generate a TAM-inducible adipocyte-specific FASN/Gsα double knockout mouse line. All of the studies performed were approved by the Institutional Animal Care and Use Committee (IACUC) of the University of Massachusetts Medical School.

### Cold Exposure

Wild type mice were single-caged and housed at 6°C for six days. Animals were exposed to a standard 12h:12h light:dark cycle and had free access to food pellets and water. Rectal temperatures were recorded every hour for a total of 6h.

### Western Blots

For protein expression analyses, iWAT was homogenized in TNET buffer (50 mM Tris-HCl, pH 7.6, 150 mM NaCl, 5 mM EDTA, 1% Triton X-100) as described^77^ with Halt protease and phosphatase inhibitors (Thermo Pierce). Samples from tissue lysates were then resolved by SDS-PAGE. Immunoblotting was performed using standard protocols. Membranes were blotted with the following antibodies: anti-FASN (BD Biosciences); anti-UCP1 (Abcam); anti-TH (Millipore); anti-phospho HSL-S660 and anti-phospho perilipin (Cell Signaling Technology), anti-Actin and anti-Tubulin (Sigma-Aldrich).

### RNA isolation and RT-qPCR

Total RNA was isolated from mouse tissues using QIAzol Lysis Reagent Protocol (QIAGEN) following the manufacturer’s instructions. cDNA was synthesized from 1μg of total RNA using iScript cDNA Synthesis Kit (BioRad). Quantitative RT-PCR was performed using iQ SybrGreen Supermix on a BioRad CFX97 thermocycler and analyzed as previously described^78,79^. *36B4, 18S* and *βM2* served as controls for normalization. Primer sequences used for qRT–PCR analyses were listed in Supplementary Table 1.

### Histological Analysis

For the immunohistochemistry (IHC) and immunofluorescence (IF) analyses, adipose tissue samples were fixed in 4% paraformaldehyde and embedded in paraffin. Sectioned slides were then stained with H&E, anti-UCP1 (Abcam, ab10983), anti-TH (Millipore, AB152) and anti-F4/80 (Bio-rad, MCA497GA) at the UMass Medical School Morphology Core. Photos from the fluorescent cells were taken with an Axiovert 35 Zeiss microscope (Zeiss) equipped with an Axiocam CCl camera at indicated magnification. For detection of macrophages in obese adipose tissue, epidydimal fat (eWAT) from 12-week-old ob/ob mice (JAX Lab) was fixed and stained with anti-F4/80 antibody as described.

### Adipo-Clear

The Adipo-Clear method was performed following the protocol described previously^11^. Mice were anesthetized, and an intracardiac perfusion/fixation was performed with 15 mL of cold 1× PBS + heparin (100 units/mL) followed by 15 mL of 4% PFA (Thermo). All harvested samples were postfixed in 4% PFA at 4°C overnight. Fixed samples were washed in 1× PBS for 1h three times at RT. The samples underwent delipidation and permeabilization following the same protocols described previously^11^. Samples were incubated in primary antibody dilutions for 4 days. In this study, anti-UCP1 (1:200, Abcam, ab10983) and anti-TH (1:200, Millipore, AB1542) were used. Secondary antibodies conjugated with Alexa-568 and Alexa-647 were purchased from Invitrogen (1:200). All whole-tissue samples were imaged on a light-sheet microscope (Ultramicroscope II; LaVision Biotec) equipped with 1.1X, 4X and 12X objective lenses and sCMOs camera (Andor Neo). Images were acquired with InspectorPro software (LaVision BioTec). 3D reconstruction images were generated using Imaris x64 software (version 8.0.1, Bitplane).

### Adipose sympathetic denervation

Adipose nerve fibers were chemically (6-hydroxydopamine, 6-OHDA) or surgically denervated as previously described^38^. Regarding 6-OHDA, a slight modification was made in the final concentration: 24μL of 6-OHDA from 100 μg/μL stock solution was administrated along each fat pad. For the cold exposures, mice were single-caged and housed at 6°C for six days following three days of post-denervation recovery. For iAdFASNKO mice, the mice were once daily treated with TAM for six consecutive days via intraperitoneal (i.p.) injection following three days of post-denervation recovery. After two weeks, animals were euthanized, and tissues were harvested. For the cold-exposed and iAdFASNKO groups, iWAT tyrosine hydroxylase content was examined via western blotting to confirm the efficiency of denervation.

### Adipose Tissue Macrophage Depletion

Macrophages were depleted from mice using clodronate-containing liposomes. iAdFASNKO mice were injected i.p. with 500 μl of clodronate (150 mg/kg/mouse) every other day for two weeks. For the control group, the same protocol was applied using the PBS control liposomes. Gene expression analysis for macrophage markers was performed to validate the macrophage depletion in the iWAT. Clodronate and PBS liposomes were provided by Liposoma BV.

### Single Cell RNA-seq of Stromal Vascular Fraction from iWAT

#### Isolation of Stromal Vascular Fraction

Whole inguinal white adipose tissue (iWAT) pads from control (n=7) and iAdFASNKO (n=8) were dissected and placed in 50 ml conical tubes containing PBS. Fat pads were cut and minced in digestion buffer (5% BSA and 2 mg/ml collagenase type II) and incubated in a 37°C water bath for 45 min. The digested tissue was passed through a 100-μm-pore-size cell strainer and centrifuged at 600g for 10 min. The pelleted cells were collected as the Stromal Vascular Fraction (SVF), and red blood cells were lysed by incubation with red blood cell lysis buffer for 5 min. The SVF was centrifuged again, and the cell pellet was resuspended in 1 ml of PBS. This final cell suspension solution was passed through a 40um cell strainer to discard debris and get single cells for Drop-Seq application.

#### Magnetic Bead Enrichment of Cellular Subtypes

SVF from control and iAdFASNKO mice were fractionated into lineage+ and lineage- pools. The SVF were labeled with microbead-tagged Direct Lineage Cell Depletion Cocktail (anti-CD5, CD11b, CD45R (B220), Anti-Gr-1 (Ly-6G/C), 7-4, and Ter-119; Miltenyi Biotec, Cat. No. 130-110-470) and passed onto MS columns (Miltenyi Biotec, Cat. No. 130-042-201) following the manufacturer’s protocol. The flow through (Lineage-) was collected and the bound cells were eluted (Lineage+ fraction). Cell fractions were resuspended in PBS with 0.04% BSA, counted, and diluted to a concentration of 1,000 cells/μL.

#### Drop-seq single cell barcoding and library preparation

Following magnetic cell sorting, single cell suspensions from lineage-positive (Lin+) and lineage-negative (Lin-), from control and iAdFASNKO were loaded onto the Single Cell 3’ Chip. Approximately 7,000 cells were loaded per channel for an expected recovery of ∼5,000 cells. The loaded Single Cell 3’ Chip was placed on a 10X Genomics Chromium Controller Instrument (10X Genomics) to generate single cell gel beads in emulsion (GEMs). Single cell RNA-seq libraries were prepared using the Chromium Single Cell 3’ GEM, Library & Gel Bead Kit v3 (10X Genomics) according to the manufacturer’s protocol.

#### Illumina high-throughput sequencing libraries

The 10X genomics library molar concentration was quantified by Qubit Fluorometric Quantitation (Thermo Fisher) and library fragment length was estimated using a TapeStation System (Agilent). Libraries were sequenced using Illumina HiSeq 3000 system performed by GENEWIZ, LLC (South Plainfield, NJ, USA) with 2.5 billion reads per run, yielding ∼625 million reads per sample.

#### Clustering for single cell RNA-seq

The Cell Ranger Single Cell Software Suite v.3.1.0 (10X Genomics) was used to perform sample de-multiplexing, alignment, filtering, and UMI counting, normalization between samples, and clustering (except for macrophage sub- clustering). Lin+ and Lin- from control and iAdFASNKO data were aggregated for direct comparison of single cell transcriptomes. Clustering and gene expression were visualized with Loupe Cell Browser v.3.1.1 (10X Genomics). Clusters were generated by Cell Ranger via K-means clustering based on the PCA of the transcriptomic signature. Clusters were manually named based on the top 20 upregulated differentially expressed genes between clusters. Related to the sub- clustering, Lin+ cells in the Macrophage-like cluster were separated into five new sub-clusters via visual inspection within the Loupe Cell Browser v.3.1.1 (10X Genomics) based on the top 20 upregulated genes for each cluster and knowledge of canonical cell markers. The heatmap was generated from the Loupe browser of the top 5 up-regulated differentially expressed genes between sub-clusters. Colors (values) are z-score of the average log fold-change between sub-clusters for each gene. To produce bar plots of the mean and variance of gene expression and violin plots of gene expression, cluster identities and filtered gene matrices generated by Cell Ranger Single Cell Software Suite v.3.1.0 (10X Genomics) were used as input into the open-source R toolkit Seurat. Cell-type pathway enrichment analysis were performed using gene ontology (GO) and KEGG pathway, related to the biological process of differential gene expression.

## Supporting information

Henriques et al - Extended Data Figures 1-5

Henriques et al - Supplementary Table 1

Up regulated genes - iAdFASNKO - Macrophage-like Sub-clustering

Down regulated genes - iAdFASNKO - Macrophage-like Sub-clustering

## Statistical analysis

Data were analyzed in GraphPad Prism 8 (GraphPad Software, Inc.). Student’s t-test was employed for the parametric data. Comparisons between more than 2 groups were performed using ANOVA with Tukey’s post-hoc test. The data are presented as means ± SEM. The significance level was set at p ≤ 0.05.

## Acknowledgments

We thank all members of the Czech Lab for helpful discussions and critical reading of the manuscript. We thank the UMass Morphology Core for assistance. This work was supported by NIH grants DK30898 and DK103047 to M.P.C. and partially supported by the Intramural Research Program of NIDDK, NIH, to L.S.W. The authors gratefully acknowledge the commitment and support by the American Diabetes Association (ADA) – Grant #1-19-PMF-035 to F.H.

## Author contributions

F.H., A.G. and M.P.C. conceived the study and designed the experiments; F.H., A.H.B., A.G., M.K., L.R. and B.Y. performed the experiments; F.H., A.H.B., M.K. performed the denervation experiments; F.H., A.H.B., L.M.F., K.B., J.C. and P.C. performed the Adipo-Clear experiments; F.H., A.H.B., L.M.F., S.K., Y.W., and J.L. performed the single cell RNA-seq data; J.D.L provided the Nrg4 whole-body knockout mice; L.S.W provided the conditional Gsα-floxed mice; F.H. and M.P.C. wrote the manuscript; F.H., A.H.B., A.G. and M.P.C. edited the final manuscript.

---

**Extended Data Movie 1**: 3D Visualization of the sympathetic innervation and browning of iWAT from control, iAdFASNKO and cold exposed mice, related to Figure 1. The movie shows rotating 3D projections of the whole iWAT depot. Anti-tyrosine hydroxylase is shown in green and UCP1 in orange. The samples were imaged at 1.1× magnification on the lightsheet microscope. Scale bars are provided for each panel.

**Extended Data Movie 2**: Low (left side) and high-resolution (right side) 3D projections of the whole iWAT after cold exposed, related to Figure 1. Anti-tyrosine hydroxylase is shown in green and UCP1 in orange. For the high-resolution (right side) imaging, the iWAT was imaged at 12X magnification on the lightsheet microscope. Scale bars are provided for each panel.

**Extended Data Movie 3**: The movie shows more details about the high-resolution 3D projections of the whole iWAT after cold exposure, related to Figure 1 and Extended Data Movie 2. Anti-tyrosine hydroxylase is shown in green and UCP1 in orange. Interestingly, using this level of magnification, it is possible to see the intimate architecture between the nerve fibers and adipocytes. Furthermore, we can observe the presence of multilocular UCP1 positive cells. Scale bars are provided for each panel.

**Extended Data Movie 4**: High-resolution (70μm) fly-through of the optical sections of iWAT from iAdFASNKO and cold exposure groups related to Figure 1. Anti-tyrosine hydroxylase is shown in green and UCP1 in orange. The samples were imaged at 24X magnification on the lightsheet microscope. Scale bars are provided for each panel.

**Extended Data Movie 5**: Ultra-resolution (10μm) fly-through of the optical sections of iWAT from iAdFASNKO and cold exposure groups related to Figure 1. Anti-tyrosine hydroxylase is shown in green and UCP1 in orange. The samples were imaged at 24X magnification on the lightsheet microscope. Scale bars are provided for each panel.

**Extended Data Figure 1: Chemical and surgical denervation fail to block upregulation of PKA signaling, TH and UCP1 expression levels in iWAT from iAdFASNKO, related to Figure 2. (a):** Depicted are representative immunoblots to detect FASN, TH, UCP1, phospho HSL, phospho perilipin, tubulin and actin protein levels in iWAT from control and iAdFASNKO mice, treated with PBS or chemically denervated with 6OHDA. **(b,c)** Confirmation that TH levels were reduced in the iWAT after surgical denervation. TH signals in iWAT were detected by immunofluorescence staining of tissue samples from **(b)** cold exposed mice and **(c)** iAdFASNKO mice. To evaluate the UCP1 protein level in the iWAT, immunohistochemistry and western blotting for UCP1 was performed for **(b,d)** cold exposure and **(c,e)** iAdFASNKO. N = 3-4 per group. These results are representative of five independent experiments.

**Extended Data Figure 2: Single Cell RNA-Seq Workflow and UMAP plot from control vs. iAdFASNKO mice, related to Figure 4. (a)** Workflow overview exhibiting all steps for the single cell RNA-seq. See the methods section for the complete details. **(b)** UMAP plot from figure 4 split into the cells from control and iAdFASNKO. The percentage of the total number of cells is shown for each cluster.

**Extended Data Figure 3: Violin plots of the top expression levels for representative genes in each different cluster, related to Figure 4**. Violin plots of log_e_ (normalized values) for the top representative genes from each cluster in the aggregated dataset. Cluster-enriched genes are shown in columns below the indicated cluster.

**Extended Data Figure 4: Representative gene for each different macrophage-like sub-cluster identified, related to Figure 4**. Log_e_ of normalized expression values for representative genes in the different Macrophage-like sub-clusters. These representative genes were used to identify and name the following clusters: Resident Macrophage (Fcgr1), M1 Macrophage (Icam2), M2 Macrophage (Mrc1), Neutrophils Cells (S100a9) and Dendritic Cells (Ccl22).

**Supplementary Figure 5: Cell proliferation analyses and immune cell infiltration, related to Figure 5. (a)** The Click-iT® EdU assay and flow cytometric analysis were conducted to measure the proliferation ratios of different types of cells present in the SVF derived from iWAT. Immune cells were defined as CD45+, progenitor cells as PDGFRα^+^ and endothelial cells as CD31+. **(b)** Immunohistochemistry for detection of F4/80 in the iWAT from wild-type mice housed at 22°C or 6°C. Red arrows indicated the presence of F4/80 positive cells in between the multilocular cells (beige cells). Bars are indicated in the panel. **(c)** Validation of the anti-F4/80 antibody. Immunofluorescent analysis to detect F4/80 signals in eWAT from ob/ob mice. Positive control for enhanced macrophage infiltration in iWAT, confirming the specificity of the anti-F4/80 antibody used. Scale bars are provided in the panel.

**Supplementary Table 1:** Primer sequences used for qRT-PCR.

## REFERENCES

1 Czech, M. P. Mechanisms of insulin resistance related to white, beige, and brown adipocytes. Mol Metab 34, 27–42, doi:10.1016/j.molmet.2019.12.014 (2020).

2 Rosen, E. D. & Spiegelman, B. M. Adipocytes as regulators of energy balance and glucose homeostasis. Nature 444, 847–853, doi:10.1038/nature05483 (2006).

3 Chouchani, E. T. & Kajimura, S. Metabolic adaptation and maladaptation in adipose tissue. Nat Metab 1, 189–200, doi:10.1038/s42255-018-0021-8 (2019).

4 Wu, J. et al. Beige adipocytes are a distinct type of thermogenic fat cell in mouse and human. Cell 150, 366–376, doi:10.1016/j.cell.2012.05.016 (2012).

5 Scheele, C. & Wolfrum, C. Brown Adipose Crosstalk in Tissue Plasticity and Human Metabolism. Endocr Rev 41, doi:10.1210/endrev/bnz007 (2020).

6 Villarroya, J. et al. New insights into the secretory functions of brown adipose tissue. J Endocrinol 243, R19–R27, doi:10.1530/JOE-19-0295 (2019).

7 Villarroya, F., Cereijo, R., Villarroya, J. & Giralt, M. Brown adipose tissue as a secretory organ. Nat Rev Endocrinol 13, 26–35, doi:10.1038/nrendo.2016.136 (2017).

8 Herz, C. T. & Kiefer, F. W. Adipose tissue browning in mice and humans. J Endocrinol 241, R97–R109, doi:10.1530/JOE-18-0598 (2019).

9 van Marken Lichtenbelt, W. D. et al. Cold-activated brown adipose tissue in healthy men. N Engl J Med 360, 1500–1508, doi:10.1056/NEJMoa0808718 (2009).

10 Saito, M. et al. High incidence of metabolically active brown adipose tissue in healthy adult humans: effects of cold exposure and adiposity. Diabetes 58, 1526–1531, doi:10.2337/db09-0530 (2009).

11 Chi, J. et al. Three-Dimensional Adipose Tissue Imaging Reveals Regional Variation in Beige Fat Biogenesis and PRDM16-Dependent Sympathetic Neurite Density. Cell Metab 27, 226–236 e223, doi:10.1016/j.cmet.2017.12.011 (2018).

12 Jiang, H., Ding, X., Cao, Y., Wang, H. & Zeng, W. Dense Intra-adipose Sympathetic Arborizations Are Essential for Cold-Induced Beiging of Mouse White Adipose Tissue. Cell Metab 26, 686–692 e683, doi:10.1016/j.cmet.2017.08.016 (2017).

13 Bartness, T. J., Vaughan, C. H. & Song, C. K. Sympathetic and sensory innervation of brown adipose tissue. Int J Obes (Lond) 34 Suppl 1, S36-42, doi:10.1038/ijo.2010.182 (2010).

14 Guilherme, A., Henriques, F., Bedard, A. H. & Czech, M. P. Molecular pathways linking adipose innervation to insulin action in obesity and diabetes mellitus. Nat Rev Endocrinol 15, 207–225, doi:10.1038/s41574-019-0165-y (2019).

15 Burl, R. B. et al. Deconstructing Adipogenesis Induced by beta3-Adrenergic Receptor Activation with Single-Cell Expression Profiling. Cell Metab 28, 300–309 e304, doi:10.1016/j.cmet.2018.05.025 (2018).

16 Weinstock, A. et al. Single-Cell RNA Sequencing of Visceral Adipose Tissue Leukocytes Reveals that Caloric Restriction Following Obesity Promotes the Accumulation of a Distinct Macrophage Population with Features of Phagocytic Cells. Immunometabolism 1, doi:10.20900/immunometab20190008 (2019).

17 Hill, D. A. et al. Distinct macrophage populations direct inflammatory versus physiological changes in adipose tissue. Proc Natl Acad Sci U S A 115, E5096–E5105, doi:10.1073/pnas.1802611115 (2018).

18 Jaitin, D. A. et al. Lipid-Associated Macrophages Control Metabolic Homeostasis in a Trem2-Dependent Manner. Cell 178, 686–698 e614, doi:10.1016/j.cell.2019.05.054 (2019).

19 Rajbhandari, P. et al. Single cell analysis reveals immune cell-adipocyte crosstalk regulating the transcription of thermogenic adipocytes. Elife 8, doi:10.7554/eLife.49501 (2019).

20 Merrick, D. et al. Identification of a mesenchymal progenitor cell hierarchy in adipose tissue. Science 364, doi:10.1126/science.aav2501 (2019).

21 Hui, X. et al. Adiponectin Enhances Cold-Induced Browning of Subcutaneous Adipose Tissue via Promoting M2 Macrophage Proliferation. Cell Metab 22, 279–290, doi:10.1016/j.cmet.2015.06.004 (2015).

22 Shan, B. et al. The metabolic ER stress sensor IRE1alpha suppresses alternative activation of macrophages and impairs energy expenditure in obesity. Nat Immunol 18, 519–529, doi:10.1038/ni.3709 (2017).

23 Lv, Y. et al. Adrenomedullin 2 Enhances Beiging in White Adipose Tissue Directly in an Adipocyte-autonomous Manner and Indirectly through Activation of M2 Macrophages. J Biol Chem 291, 23390–23402, doi:10.1074/jbc.M116.735563 (2016).

24 Ceddia, R. P. & Collins, S. A compendium of G-protein-coupled receptors and cyclic nucleotide regulation of adipose tissue metabolism and energy expenditure. Clin Sci (Lond) 134, 473–512, doi:10.1042/CS20190579 (2020).

25 Li, Y. Q. et al. Gsalpha deficiency in adipose tissue improves glucose metabolism and insulin sensitivity without an effect on body weight. Proc Natl Acad Sci U S A 113, 446–451, doi:10.1073/pnas.1517142113 (2016).

26 Harris, R. B. S. Denervation as a tool for testing sympathetic control of white adipose tissue. Physiol Behav 190, 3–10, doi:10.1016/j.physbeh.2017.07.008 (2018).

27 Blaszkiewicz, M., Willows, J. W., Johnson, C. P. & Townsend, K. L. The Importance of Peripheral Nerves in Adipose Tissue for the Regulation of Energy Balance. Biology (Basel) 8, doi:10.3390/biology8010010 (2019).

28 Collins, S. beta-Adrenoceptor Signaling Networks in Adipocytes for Recruiting Stored Fat and Energy Expenditure. Front Endocrinol (Lausanne) 2, 102, doi:10.3389/fendo.2011.00102 (2011).

29 Li, G. et al. Intermittent Fasting Promotes White Adipose Browning and Decreases Obesity by Shaping the Gut Microbiota. Cell Metab 26, 672–685 e674, doi:10.1016/j.cmet.2017.08.019 (2017).

30 Fabbiano, S. et al. Caloric Restriction Leads to Browning of White Adipose Tissue through Type 2 Immune Signaling. Cell Metab 24, 434–446, doi:10.1016/j.cmet.2016.07.023 (2016).

31 Aldiss, P. et al. Exercise-induced ‘browning’ of adipose tissues. Metabolism 81, 63–70, doi:10.1016/j.metabol.2017.11.009 (2018).

32 Patsouris, D. et al. Burn Induces Browning of the Subcutaneous White Adipose Tissue in Mice and Humans. Cell Rep 13, 1538–1544, doi:10.1016/j.celrep.2015.10.028 (2015).

33 Liu, D. et al. Activation of mTORC1 is essential for beta-adrenergic stimulation of adipose browning. J Clin Invest 126, 1704–1716, doi:10.1172/JCI83532 (2016).

34 Guilherme, A. et al. Adipocyte lipid synthesis coupled to neuronal control of thermogenic programming. Mol Metab 6, 781–796, doi:10.1016/j.molmet.2017.05.012 (2017).

35 Guilherme, A. et al. Neuronal modulation of brown adipose activity through perturbation of white adipocyte lipogenesis. Mol Metab 16, 116–125, doi:10.1016/j.molmet.2018.06.014 (2018).

36 Lodhi, I. J. et al. Inhibiting adipose tissue lipogenesis reprograms thermogenesis and PPARgamma activation to decrease diet-induced obesity. Cell Metab 16, 189–201, doi:10.1016/j.cmet.2012.06.013 (2012).

37 Cao, Q., Jing, J., Cui, X., Shi, H. & Xue, B. Sympathetic nerve innervation is required for beigeing in white fat. Physiol Rep 7, e14031, doi:10.14814/phy2.14031 (2019).

38 Vaughan, C. H., Zarebidaki, E., Ehlen, J. C. & Bartness, T. J. Analysis and measurement of the sympathetic and sensory innervation of white and brown adipose tissue. Methods Enzymol 537, 199–225, doi:10.1016/B978-0-12-411619-1.00011-2 (2014).

39 Harris, R. B. Sympathetic denervation of one white fat depot changes norepinephrine content and turnover in intact white and brown fat depots. Obesity (Silver Spring) 20, 1355–1364, doi:10.1038/oby.2012.95 (2012).

40 Shi, H., Song, C. K., Giordano, A., Cinti, S. & Bartness, T. J. Sensory or sympathetic white adipose tissue denervation differentially affects depot growth and cellularity. Am J Physiol Regul Integr Comp Physiol 288, R1028–1037, doi:10.1152/ajpregu.00648.2004 (2005).

41 Dickson, L. M., Gandhi, S., Layden, B. T., Cohen, R. N. & Wicksteed, B. Protein kinase A induces UCP1 expression in specific adipose depots to increase energy expenditure and improve metabolic health. Am J Physiol Regul Integr Comp Physiol 311, R79–88, doi:10.1152/ajpregu.00114.2016 (2016).

42 Garretson, J. T. et al. Lipolysis sensation by white fat afferent nerves triggers brown fat thermogenesis. Mol Metab 5, 626–634, doi:10.1016/j.molmet.2016.06.013 (2016).

43 Foster, M. T. & Bartness, T. J. Sympathetic but not sensory denervation stimulates white adipocyte proliferation. Am J Physiol Regul Integr Comp Physiol 291, R1630–1637, doi:10.1152/ajpregu.00197.2006 (2006).

44 Wang, G. X. et al. The brown fat-enriched secreted factor Nrg4 preserves metabolic homeostasis through attenuation of hepatic lipogenesis. Nat Med 20, 1436–1443, doi:10.1038/nm.3713 (2014).

45 Comas, F. et al. Neuregulin 4 Is a Novel Marker of Beige Adipocyte Precursor Cells in Human Adipose Tissue. Front Physiol 10, 39, doi:10.3389/fphys.2019.00039 (2019).

46 Pfeifer, A. NRG4: an endocrine link between brown adipose tissue and liver. Cell Metab 21, 13–14, doi:10.1016/j.cmet.2014.12.008 (2015).

47 Bluher, M. Neuregulin 4: A “Hotline” Between Brown Fat and Liver. Obesity (Silver Spring) 27, 1555–1557, doi:10.1002/oby.22595 (2019).

48 Rosell, M. et al. Brown and white adipose tissues: intrinsic differences in gene expression and response to cold exposure in mice. Am J Physiol Endocrinol Metab 306, E945–964, doi:10.1152/ajpendo.00473.2013 (2014).

49 Pellegrinelli, V. et al. Adipocyte-secreted BMP8b mediates adrenergic-induced remodeling of the neuro-vascular network in adipose tissue. Nat Commun 9, 4974, doi:10.1038/s41467-018-07453-x (2018).

50 Rondini, E. A. & Granneman, J. G. Single cell approaches to address adipose tissue stromal cell heterogeneity. Biochem J 477, 583–600, doi:10.1042/BCJ20190467 (2020).

51 Cereijo, R. et al. CXCL14, a Brown Adipokine that Mediates Brown-Fat-to-Macrophage Communication in Thermogenic Adaptation. Cell Metab 28, 750–763 e756, doi:10.1016/j.cmet.2018.07.015 (2018).

52 Jun, H. et al. An immune-beige adipocyte communication via nicotinic acetylcholine receptor signaling. Nat Med 24, 814–822, doi:10.1038/s41591-018-0032-8 (2018).

53 Pirzgalska, R. M. et al. Sympathetic neuron-associated macrophages contribute to obesity by importing and metabolizing norepinephrine. Nat Med 23, 1309–1318, doi:10.1038/nm.4422 (2017).

54 Villarroya, F., Cereijo, R., Villarroya, J., Gavalda-Navarro, A. & Giralt, M. Toward an Understanding of How Immune Cells Control Brown and Beige Adipobiology. Cell Metab 27, 954–961, doi:10.1016/j.cmet.2018.04.006 (2018).

55 Camell, C. D. et al. Inflammasome-driven catecholamine catabolism in macrophages blunts lipolysis during ageing. Nature 550, 119–123, doi:10.1038/nature24022 (2017).

56 Suarez-Zamorano, N. et al. Microbiota depletion promotes browning of white adipose tissue and reduces obesity. Nat Med 21, 1497–1501, doi:10.1038/nm.3994 (2015).

57 Liu, P. S., Lin, Y. W., Burton, F. H. & Wei, L. N. Injecting engineered anti-inflammatory macrophages therapeutically induces white adipose tissue browning and improves diet-induced insulin resistance. Adipocyte 4, 123–128, doi:10.4161/21623945.2014.981438 (2015).

58 Vargovic, P., Manz, G. & Kvetnansky, R. Continuous cold exposure induces an anti-inflammatory response in mesenteric adipose tissue associated with catecholamine production and thermogenin expression in rats. Endocr Regul 50, 137–144, doi:10.1515/enr-2016-0015 (2016).

59 Marro, B. S., Legrain, S., Ware, B. C. & Oldstone, M. B. Macrophage IFN-I signaling promotes autoreactive T cell infiltration into islets in type 1 diabetes model. JCI Insight 4, doi:10.1172/jci.insight.125067 (2019).

60 van Rooijen, N. & Hendrikx, E. Liposomes for specific depletion of macrophages from organs and tissues. Methods Mol Biol 605, 189–203, doi:10.1007/978-1-60327-360-2_13 (2010).

61 Van Rooijen, N. & Sanders, A. Liposome mediated depletion of macrophages: mechanism of action, preparation of liposomes and applications. J Immunol Methods 174, 83–93, doi:10.1016/0022-1759(94)90012-4 (1994).

62 Zeisberger, S. M. et al. Clodronate-liposome-mediated depletion of tumour-associated macrophages: a new and highly effective antiangiogenic therapy approach. Br J Cancer 95, 272–281, doi:10.1038/sj.bjc.6603240 (2006).

63 Amano, S. U. et al. Local proliferation of macrophages contributes to obesity-associated adipose tissue inflammation. Cell Metab 19, 162–171, doi:10.1016/j.cmet.2013.11.017 (2014).

64 Bu, L., Gao, M., Qu, S. & Liu, D. Intraperitoneal injection of clodronate liposomes eliminates visceral adipose macrophages and blocks high-fat diet-induced weight gain and development of insulin resistance. AAPS J 15, 1001–1011, doi:10.1208/s12248-013-9501-7 (2013).

65 Feng, B. et al. Clodronate liposomes improve metabolic profile and reduce visceral adipose macrophage content in diet-induced obese mice. PLoS One 6, e24358, doi:10.1371/journal.pone.0024358 (2011).

66 Reverte-Salisa, L., Sanyal, A. & Pfeifer, A. Role of cAMP and cGMP Signaling in Brown Fat. Handb Exp Pharmacol 251, 161–182, doi:10.1007/164_2018_117 (2019).

67 Cao, Y., Wang, H. & Zeng, W. Whole-tissue 3D imaging reveals intra-adipose sympathetic plasticity regulated by NGF-TrkA signal in cold-induced beiging. Protein Cell 9, 527–539, doi:10.1007/s13238-018-0528-5 (2018).

68 Nguyen, K. D. et al. Alternatively activated macrophages produce catecholamines to sustain adaptive thermogenesis. Nature 480, 104–108, doi:10.1038/nature10653 (2011).

69 Qiu, Y. et al. Eosinophils and type 2 cytokine signaling in macrophages orchestrate development of functional beige fat. Cell 157, 1292–1308, doi:10.1016/j.cell.2014.03.066 (2014).

70 Fischer, K. et al. Alternatively activated macrophages do not synthesize catecholamines or contribute to adipose tissue adaptive thermogenesis. Nat Med 23, 623–630, doi:10.1038/nm.4316 (2017).

71 Yang, H. & Yang, L. Targeting cAMP/PKA pathway for glycemic control and type 2 diabetes therapy. J Mol Endocrinol 57, R93–R108, doi:10.1530/JME-15-0316 (2016).

72 Braun, K., Oeckl, J., Westermeier, J., Li, Y. & Klingenspor, M. Non-adrenergic control of lipolysis and thermogenesis in adipose tissues. J Exp Biol 221, doi:10.1242/jeb.165381 (2018).

73 Gerhart-Hines, Z. et al. The cAMP/PKA pathway rapidly activates SIRT1 to promote fatty acid oxidation independently of changes in NAD(+). Mol Cell 44, 851–863, doi:10.1016/j.molcel.2011.12.005 (2011).

74 Kumar, N., Liu, D., Wang, H., Robidoux, J. & Collins, S. Orphan nuclear receptor NOR-1 enhances 3’,5’-cyclic adenosine 5’-monophosphate-dependent uncoupling protein-1 gene transcription. Mol Endocrinol 22, 1057–1064, doi:10.1210/me.2007-0464 (2008).

75 Jeon, E. J., Kim, D. Y., Lee, N. H., Choi, H. E. & Cheon, H. G. Telmisartan induces browning of fully differentiated white adipocytes via M2 macrophage polarization. Sci Rep 9, 1236, doi:10.1038/s41598-018-38399-1 (2019).

76 Zhao, H. et al. Exosomes From Adipose-Derived Stem Cells Attenuate Adipose Inflammation and Obesity Through Polarizing M2 Macrophages and Beiging in White Adipose Tissue. Diabetes 67, 235–247, doi:10.2337/db17-0356 (2018).

77 Morley, T. S., Xia, J. Y. & Scherer, P. E. Selective enhancement of insulin sensitivity in the mature adipocyte is sufficient for systemic metabolic improvements. Nat Commun 6, 7906, doi:10.1038/ncomms8906 (2015).

78 Livak, K. J. & Schmittgen, T. D. Analysis of relative gene expression data using real-time quantitative PCR and the 2(-Delta Delta C(T)) Method. Methods 25, 402–408, doi:10.1006/meth.2001.1262 (2001).

79 Schmittgen, T. D. & Livak, K. J. Analyzing real-time PCR data by the comparative C(T) method. Nat Protoc 3, 1101–1108 (2008).

