## Supplementary figures and images for "Single Cell RNA Profiling Reveals Adipocyte to Macrophage Signaling Sufficient to Enhance Thermogenesis"

### Henriques et al - Extended Data Figures 1-5

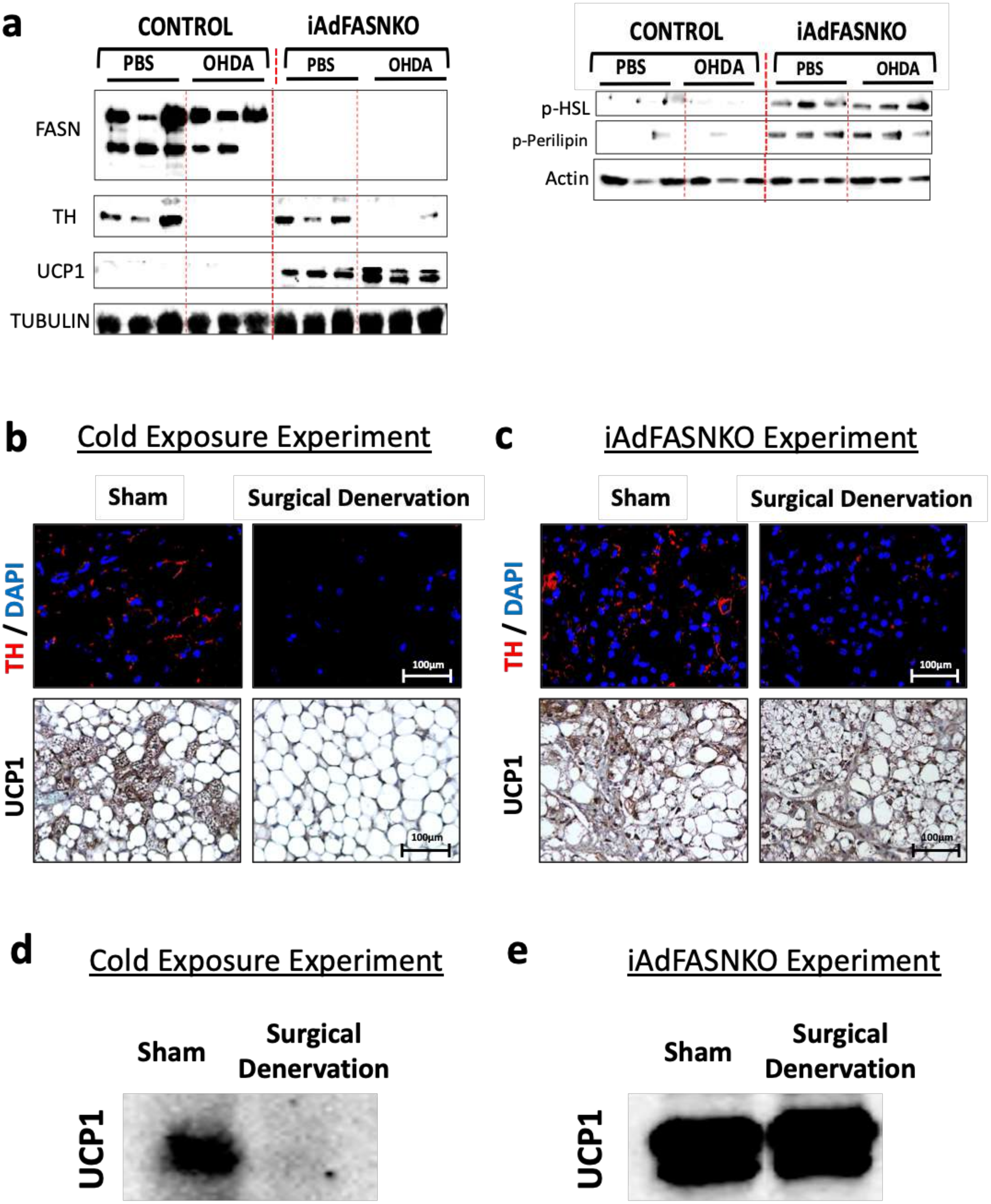


**Extended Data Fig. 1**

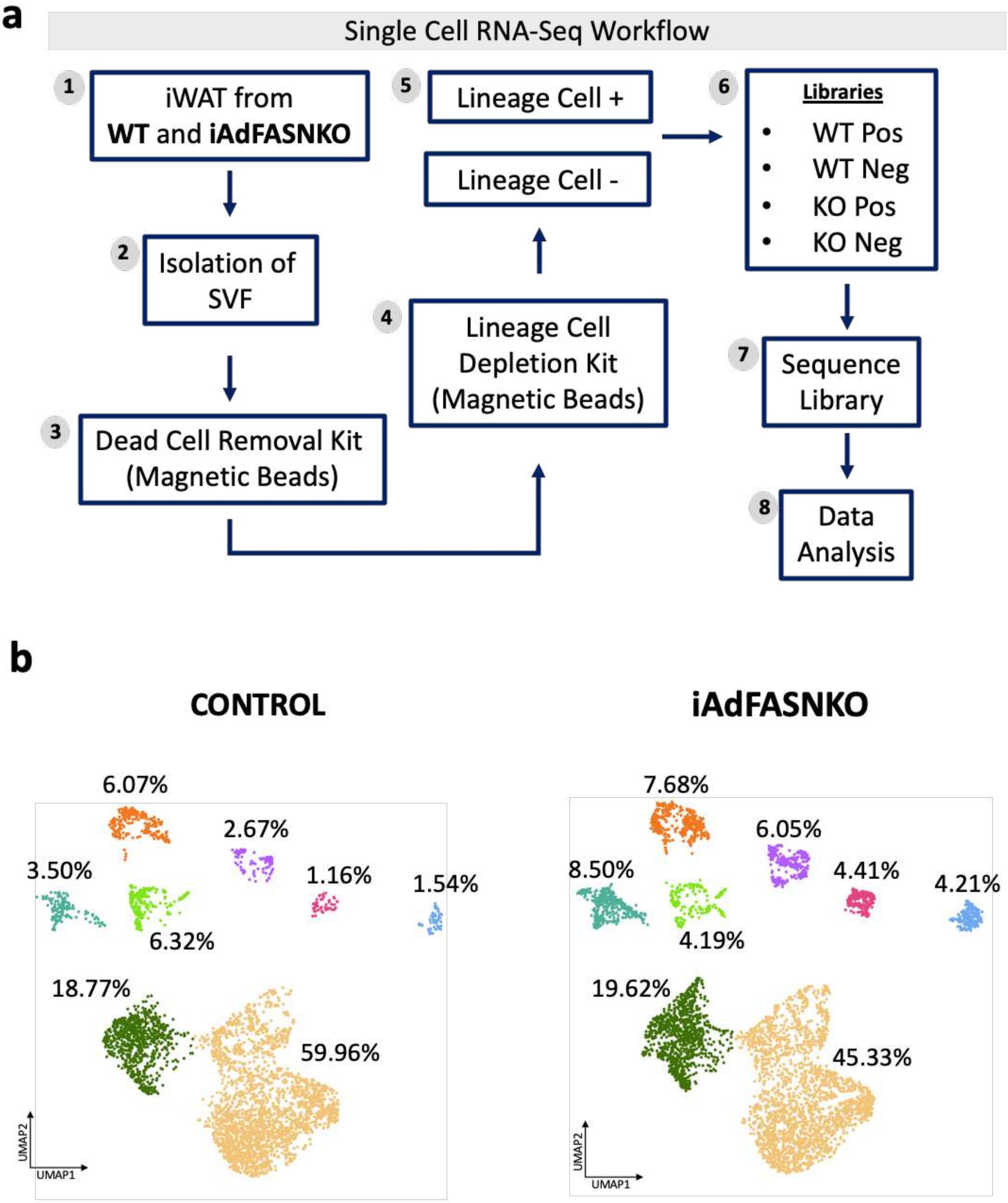


**Extended Data Fig. 2**

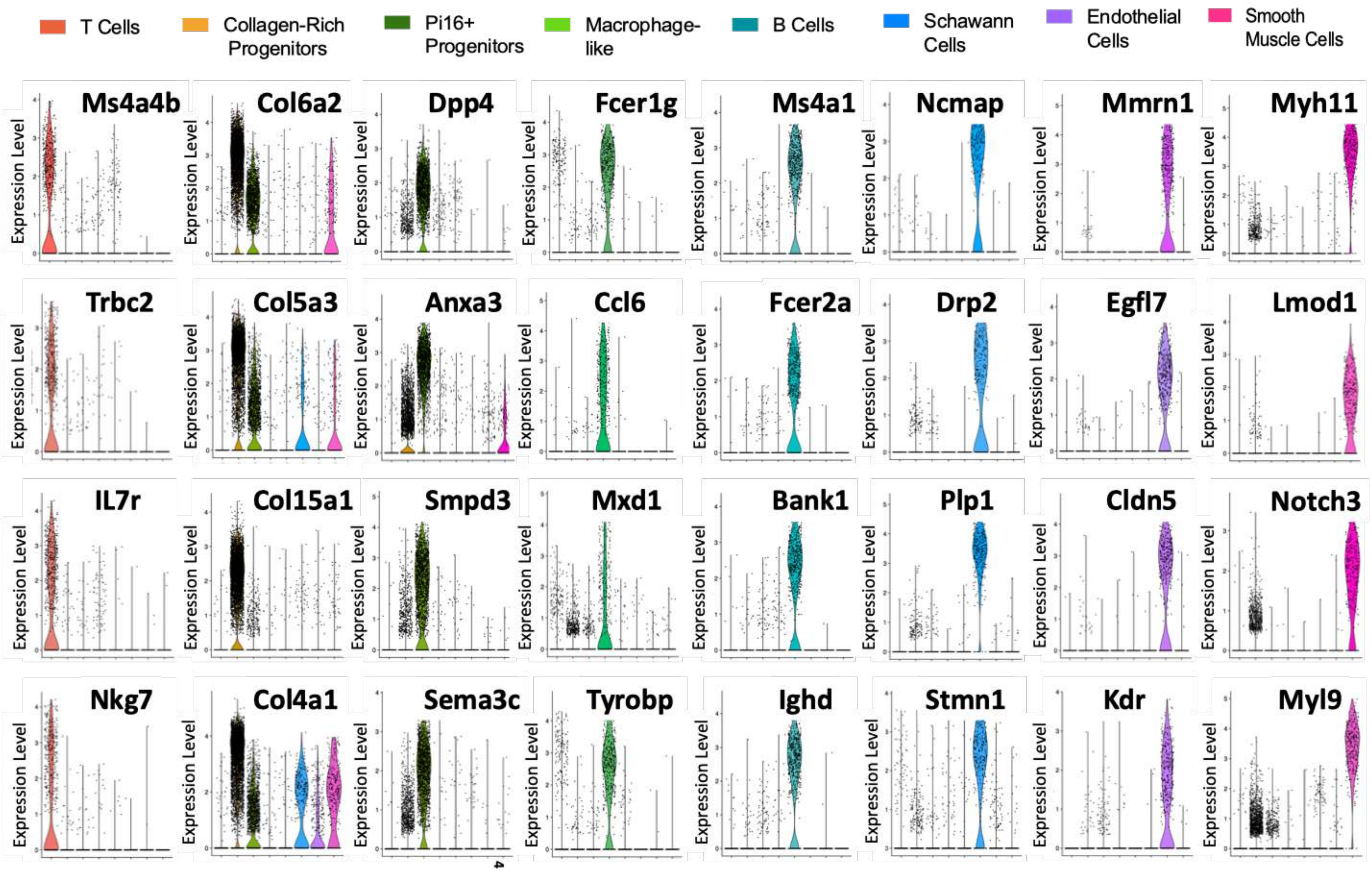


**Extended Data Fig. 3**

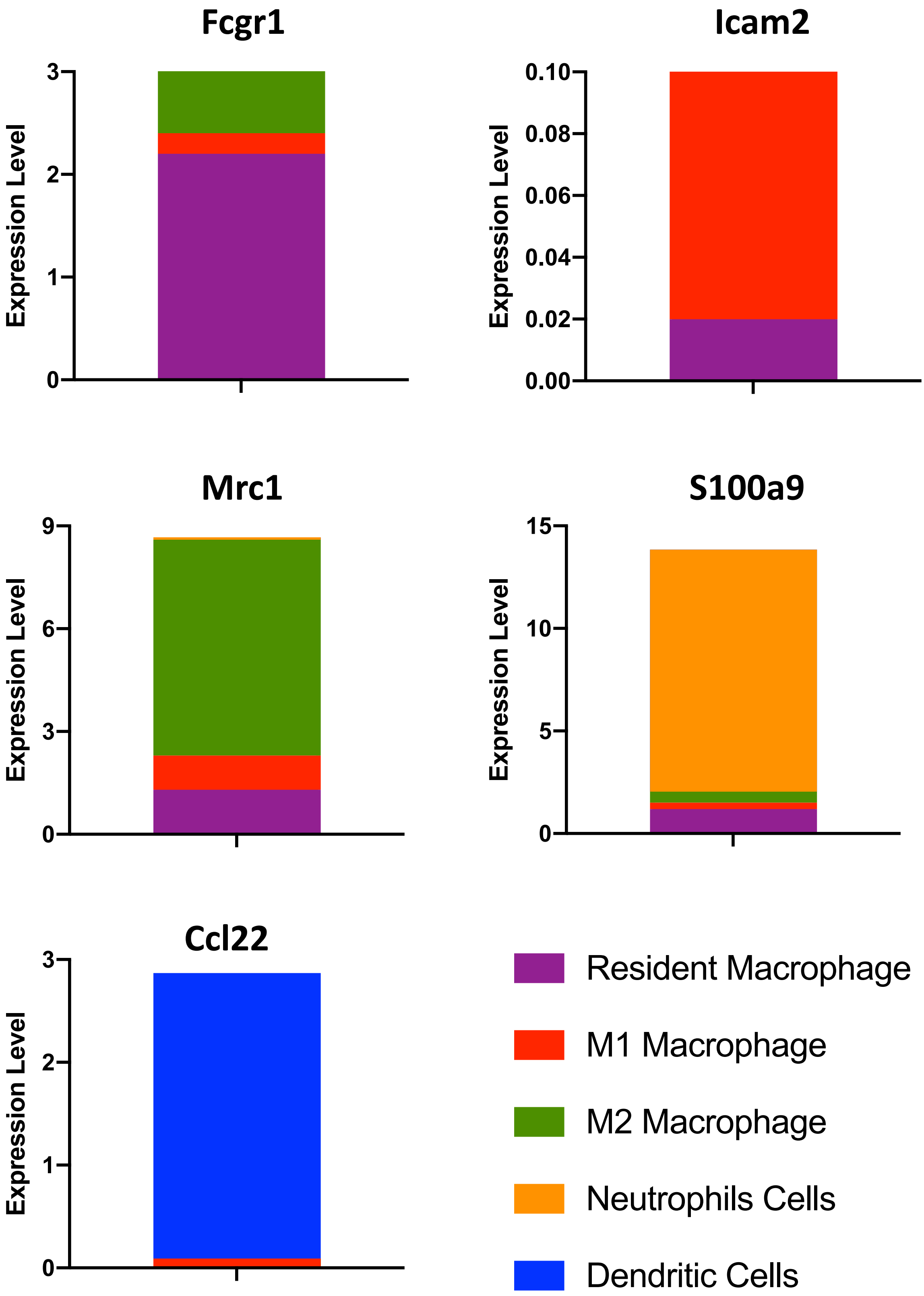


**Extended Data Fig. 4**

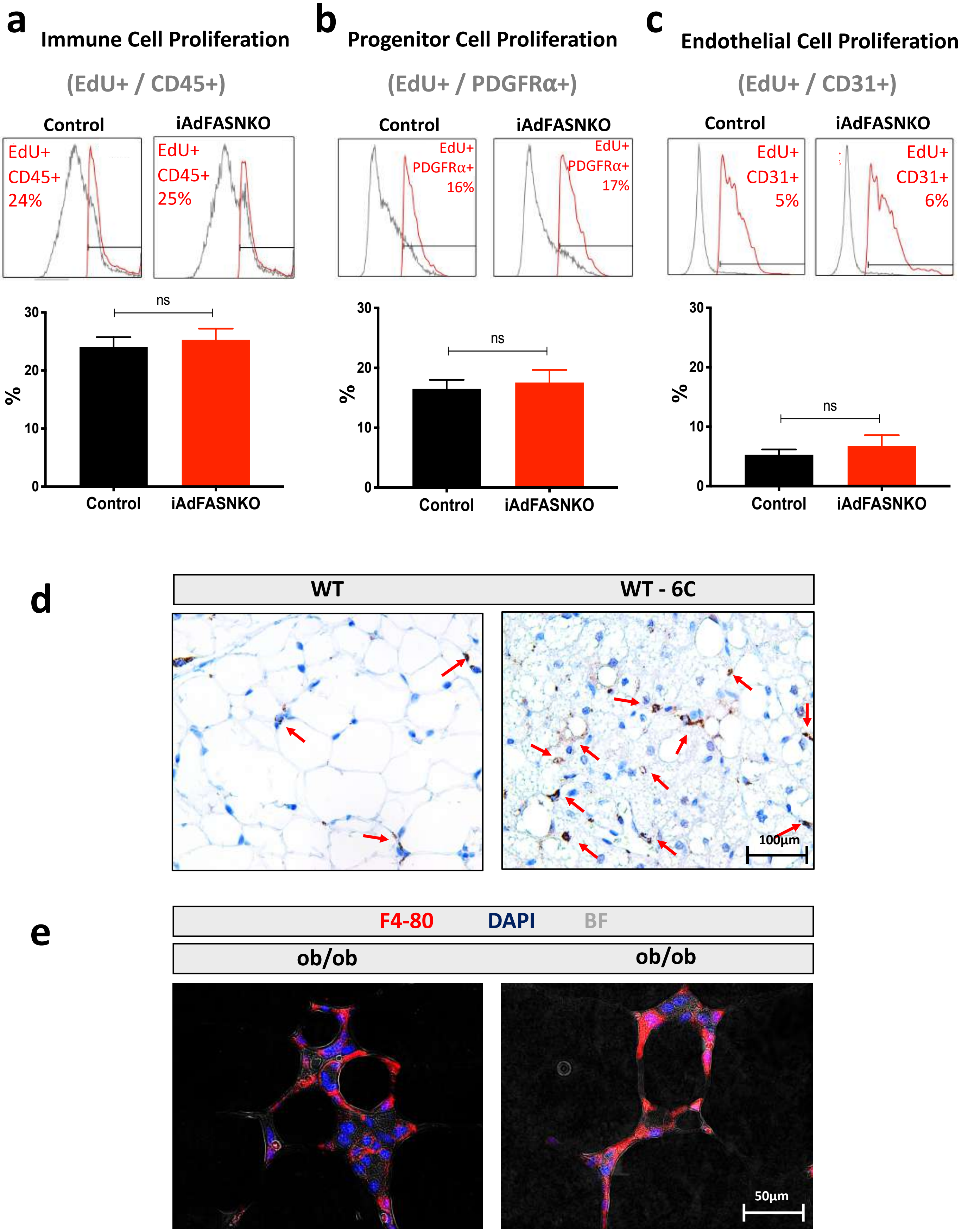


Extended Data Fig. 5
