## Supplementary material for "Single Cell RNA Profiling Reveals Adipocyte to Macrophage Signaling Sufficient to Enhance Thermogenesis": Henriques et al - Supplementary Table 1

| GENE | Forward | Reverse |
|---|---|---|
| Ucp1 | ACTGCCACACCTCCAGTCATT | CTTTGCCTCACTCAGGATTGG |
| Fasn | GGAGGTGGTGATAGCCGGTAT | TGGGTAATCCATAGAGCCCAG |
| Cidea | ATCACAACTGGCCTGGTTACG | TACTACCCGGTGTCCATTTCT |
| Gsα | ACAAGCAGGTCTACCGGGCC | CTCCGTTAAACCCATTAACATGCA |
| Nrg4 | CACGCTGCGAAGAGGTTTTTC | CGCGATGGTAAGAGTGAGGA |
| F4/80 | CTTTGGCTATGGGCTTCCAGTC | GCAAGGAGGACAGAGTTTATCGTG |
| CD206 | CTCTGTTCAGCTATTGGACGC | TGGCACTCCCAAACATAATTTGA |
| CD11c | CTGGATAGCCTTTCTTCTGCTG | GCACACTGTGTCCGAACTCA |
| CD68 | TGTCTGATCTTGCTAGGACCG | GAGAGTAACGGCCTTTTTGTGA |
| 36B4 | TCCAGGCTTTGGGCATCA | CTTTATCAGCTGCACATCACTCAGA |
| β*2m* | CATGGCTCGCTCGGTGAC | CAGTTCAGTATGTTCGGCTTCC |
| *18S* | CGAACGTCTGCCCTATCAACTT | CCGGAATCGAACCCTGATT |

**Supplementary Table 1**
